# Host iron deficiency protects against *Plasmodium* infection and drives parasite molecular reprofiling

**DOI:** 10.1101/2025.06.24.660981

**Authors:** Danielle Clucas, Cavan Bennett, Rebecca Harding, Anne Pettikiriarachchi, Andrew Baldi, Louise M. Randall, Ryan Steel, Ronan Mellin, Melissa Hobbs, Sabrina Caiazzo, Martin N. Mwangi, Katherine L. Fielding, Peter F. Hickey, Tracey M. Baldwin, Daniela Amann-Zalcenstein, Samantha J. Emery-Corbin, Glory Mzembe, Ernest Moya, Sabine Braat, Aaron Jex, Ayse Y Demir, Hans Verhoef, Kamija S. Phiri, Beverley-Ann Biggs, Wai-Hong Tham, Justin A. Boddey, Sant-Rayn Pasricha, Ricardo Ataíde

## Abstract

Iron deficiency, anemia and Plasmodium infection represent significant global health challenges with overlapping geographical distributions, particularly affecting pregnant women in Africa, yet the mechanisms underlying their interactions remain poorly understood. We employed a multilayered approach combining clinical data from Malawian pregnant women (n=711) in the REVAMP trial, a genetic mouse model (Tmprss6-knockout), and in vitro P. falciparum cultures to clarify associations between iron status and malaria susceptibility. Iron deficiency was associated with 50% reduced P. falciparum parasitemia in pregnant women (95% CI [30%-64%], p<0.0001), while iron-deficient mice exhibited significantly improved survival against P. berghei (median 15.5 days vs. 7.0 days for WT mice) and protection from cerebral malaria (83% vs 17% survival). Iron chelation induced substantial transcriptomic and proteomic changes in cultured parasites, affecting host cell invasion and nutrient acquisition processes. Importantly, intravenous iron supplementation did not increase subsequent parasitemia when coupled with malaria prevention. These findings demonstrate that iron deficiency protects against Plasmodium infection and support WHO recommendations for iron supplementation in malaria-endemic regions when combined with adequate malaria prevention strategies.

## Introduction

Iron deficiency anemia and *Plasmodium* infection remain critical global health problems with significant overlap in their geographical distribution. In 2021, an estimated 1.92 billion people worldwide experienced anemia, while approximately 247 million cases of malaria and 619,000 malaria-related deaths were reported in the same year.^1,2^ Across much of sub-Saharan Africa, both iron deficiency and *Plasmodium* infection coexist as significant public health challenges, with anemia commonly resulting from either condition.^3–5^

These conditions appear to interact in complex ways that complicate public health interventions. There are concerns that iron supplementation might increase susceptibility to *Plasmodium* infection. These concerns stem from evidence such as a large randomized controlled trial (RCT) of ∼24,000 Tanzanian children in a malaria-endemic area, which demonstrated an increased risk of hospitalization, death, and malaria-specific hospitalization in children receiving oral iron compared to those receiving placebo.^6^ In addition, observational evidence suggests that iron deficiency may protect against malaria in children. For example, in a cohort of 785 Tanzanian children followed for up to three years, iron-deficient children had lower odds of subsequent parasitemia and reduced malaria-associated mortality.^7^

The biological mechanisms underlying these interactions are multifaceted. *Plasmodium* infection may induce iron deficiency through several pathways, including hemolysis,^8^ infection-induced hepcidin upregulation^9^, and impaired iron absorption, while, simultaneously, iron deficiency may modify host susceptibility to the parasite.^10–12^ *In vitro* studies have demonstrated reduced *P. falciparum* growth and invasion in red blood cells from anemic participants — a protective effect that was reversed when iron supplementation was administered.^8,13–15^

Pregnant women represent a particularly vulnerable population for both iron deficiency and *Plasmodium* infection.^16,17^ Approximately one-third of pregnancies in moderate and high malaria transmission countries in the African region are exposed to malaria. A similar proportion of pregnant women globally experience anemia,^18^ with half of these anemia cases expected to be iron-responsive.^19^ Both anemia and malaria during pregnancy are associated with adverse maternal and neonatal outcomes, including low birth weight (LBW), which increases the risk of neonatal and infant mortality.^17,20–23^ Importantly, iron supplementation has been linked to increases in birth weight.^24^ These complex risk-benefit considerations underlie the World Health Organization’s (WHO) current recommendation for universal iron supplementation for pregnant women in areas where anemia prevalence exceeds 40%, many of which are also malaria-endemic.^25^

Despite these recommendations, significant knowledge gaps remain regarding the safety and efficacy of iron interventions in malaria-endemic regions, particularly for pregnant women. Previous studies have been limited by challenges in accurately diagnosing iron deficiency during inflammation, confounding by malaria prevention measures, and difficulty isolating iron status’ specific effects on *Plasmodium* infection from other factors. ^26–28^

In this study, we employed a multi-system approach to clarify the association between iron deficiency and risk of malaria infection. First, we examined clinical data from the REVAMP clinical trial^29^ in Malawian anemic pregnant women to assess the relationship between iron status and *P. falciparum* parasitemia detected by ultra-sensitive PCR. Second, we utilized genetic mouse model to isolate the effect of iron deficiency on the progression of *Plasmodium* infection. Finally, we explored the direct effects of iron chelation on *P. falciparum* parasites *in vitro* through transcriptomic and proteomic analyses. This comprehensive approach allowed us to test our hypothesis that iron deficiency confers protection against *Plasmodium* infection through direct effects on parasite metabolism, which would have implications for the safety of iron supplementation in malaria-endemic regions.

## Methods

### Study design and participants

Samples were derived from the REVAMP randomized controlled trial (ANZCTR registration number ACTRN12618001268235).^29^ REVAMP was an open-label, randomized controlled trial comparing a single dose of intravenous ferric carboxymaltose at enrolment to standard-of-care oral iron for the duration of pregnancy for anemia recovery in 862 anemic (capillary hemoglobin <10.0 g/dL) second-trimester pregnant women in Blantyre and Zomba districts in Malawi.^29^ All eligible pregnant women received insecticide treated nets, and intermittent preventative treatment with sulfadoxine/pyrimethamine (IPTp-SP, standard malaria prophylaxis during pregnancy) as per the Malawian national guidelines— HIV-positive women received cotrimoxazole.²¹ The trial was approved by ethics committees at the College of Medicine, University of Malawi, Zomba, Malawi; and The Walter and Eliza Hall Institute of Medical Research, Melbourne, Victoria, Australia.

### Procedures

Before enrolment in REVAMP (i.e., pre-screening), women were screened for parasitemia by rapid diagnostic test (RDT) (SD Bioline Malaria AG P.F/PAN, Standard Diagnostics, Inc.). RDT-positive women were excluded from enrolment and were treated with artemether-lumefantrine (AL); however, if they met all other inclusion criteria, they could be rescreened after one week and enter the trial if they were malaria microscopy-negative (Figure 1a). Venous blood was collected at enrolment (baseline, prior to randomization and treatment administration), at 28 days post-treatment, 36 weeks gestation, delivery, 28 days postpartum, and unscheduled visits when participants presented with illness during the trial.^30^ EDTA whole blood was banked for *P. falciparum* DNA analysis.

**Figure 1.**
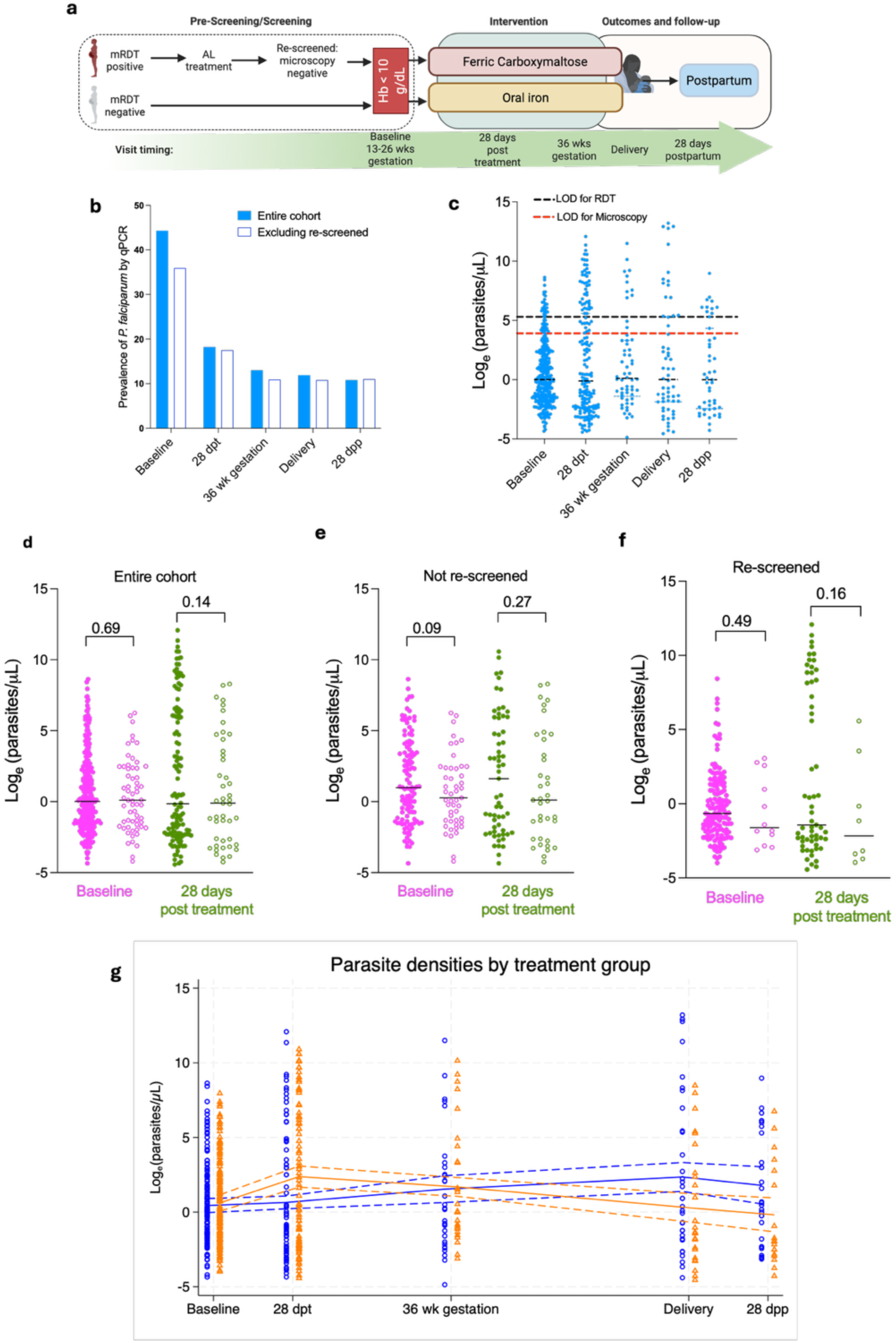
Trial cohort and *P. falciparum prevalence and parasite density.* (a) Flowchart depicting participant enrolment and sample collection timeline in the REVAMP trial. Women who tested positive by malaria rapid diagnostic test (RDT) at pre-screening received artemether-lumefantrine treatment and were re-screened after at least one week before enrolment. Created with Biorender.com (b) Prevalence of *P. falciparum* infection by ultrasensitive qPCR across study timepoints. All women (blue) or excluding those who were malaria RDT-positive at screening, treated with AL and enrolled after microscopy-negative slide (re-screened) (white). (c) Parasite densities (parasites/μL) in qPCR-positive samples across study timepoints. Box plots show median, interquartile range, and range. The theoretical limits of detection (LOD) for RDT (black) and microscopy (red) are shown on the graph. (d) Parasite densities stratified by iron status (iron-replete (IR) – full circles; iron-deficient (ID) – empty circles) at baseline (magenta) and 28 days post-treatment (green) for all women. (e) Parasite densities stratified by iron status (iron-replete (IR) – full circles; iron-deficient (ID) – empty circles) at baseline (magenta) and 28 days post-treatment (green) excluding those who were malaria RDT-positive at screening, treated with AL and enrolled after microscopy-negative slide (re-screened). (f) Parasite densities stratified by iron status (iron-replete (IR) – full circles; iron-deficient (ID) – empty circles) at baseline (magenta) and 28 days post-treatment (green) in those who were re-screened. (g) Parasite densities stratified by treatment group (IV iron (orange) vs. oral iron (blue)) across timepoints. Lines represent median values with 95% confidence intervals. dpt – days post treatment; wk – weeks; dpp – days postpartum. P-values represent two-sample t-tests (d) and (e) or Mann-Whitney (f).

#### Clinical laboratory measurements

Hemoglobin (Hb) concentration was measured on venous blood using a Sysmex XP-300 automated analyzer (Sysmex, Japan). Serum ferritin and C-reactive protein (CRP) were measured at the Meander Medical Centre laboratory (accreditation number M040, EN ISO 15189:2012, Amersfoort, The Netherlands) using the Architect System (Abbott Ireland, Longford, Ireland). The lower limit of detection for ferritin was 3.1 µg/L, and for CRP was 0.2 mg/L.

### Parasitemia detection

DNA was extracted from 400μL of EDTA whole blood using the QIAamp 96 Blood kit (Qiagen). DNA was also extracted from 200μL of reconstituted WHO International Standard for *Plasmodium falciparum* DNA Nucleic Acid Amplification Techniques (NIBSC code: 04/176, National Institute for Biological Standards and Control, UK).^31^ Pf-VarATS ultra-sensitive PCR ^32^ was performed using published probe and primer sequences and Qiagen QuantiNova Probe PCR master mix, with master mix preparation and cycling conditions as per manufacturer recommendations. The lower limit of detection was 0.12 parasites/μL of blood. qPCR was performed on the LightCycler 480 II thermocycler (Roche) with samples run in triplicate.

### Pre-clinical models of iron deficiency

#### Animals and Ethics

The iron deficiency model comprised *Tmprss6-*knockout (KO) mice on a C57Bl/6 background^33^. *Tmprss6*-heterozygous (Het) littermates and wild-type (WT) animals were used as controls. All animal experiments were conducted in accordance with the recommendations in the National Statement on Ethical Conduct in Animal Research of the National Health and Medical Research Council and under approved requirements set out by the Walter and Eliza Hall Institute of Medical Research (WEHI) Animal Ethics Committee, Melbourne, Australia (approvals 2017.031, 2019.013, 2021.064, and 2020.034). Animals were housed in specific pathogen-free conditions with access to standard chow (180mg/kg iron) and water *ad libitum*.

#### Mosquito colony maintenance

*Anopheles stephensi* mosquitoes originally imported from Johns Hopkins School of Public Health, USA, were reared and maintained in the insectary at the WEHI, Melbourne, Australia according to standard methods. The mosquitoes were stored in BugDorm insect cages as mixed genders with an environment of 26-27 °C, relative humidity 70%–80%, light/dark photoperiod 12 h:12 h including 30 min ramping to imitate dawn and dusk. Mosquitoes were fed on reverse osmosis filtered water via cotton wicks and sugar cubes (sucrose-CSR).

#### P. berghei sporozoite production

To produce *P. berghei* sporozoites expressing mCherry and luciferase reporters (PbmcherryLuci)^34^, BALB/c ‘donor’ mice were infected via the intraperitoneal (i.p.) route with blood-stage parasites and 4 days later infected erythrocytes from the donor mice were transferred to naïve “acceptor” mice via i.p. or i.v. injection. Mice with ≥1% parasitemia and exhibiting exflagellation of microgametes by microscopy at 40x magnification were anesthetized with ketamine/xylazine via i.p. inoculation and individually placed on top of a single container of 50 female *An. stephensi* (3–5 days old) mosquitoes, allowing them to feed on mice for 15-30 min, after which any unfed mosquitoes were collected and discarded. Infected mosquitoes were maintained at 21 °C with 80% humidity and 12 h:12 h light:dark photoperiod with 30 min ramping. Midgut oocysts were determined at 14 days post□blood feeding and at 18 days post□blood feeding salivary glands were dissected from cold-anesthetized and ethanol killed mosquitoes into Schneider’s Insect Media (Sigma-Aldrich) pH 7.0^35^ and quantified in a hemocytometer counting chamber (Assistent, Neubauer improved) under phase at 400x magnification.

#### P. berghei infection and IVIS imaging

Mice were infected by intravenous injection of 6500–10000 PbmCherryLuci sporozoites (Day 0). Mice were euthanized upon reaching any one of the following endpoints: signs of cerebral malaria (loss of self-righting reflex and hind-limb paresis); >20% parasitemia; or at least two of the following: weight loss >10%, hunched posture, piloerection, blanching of the tail and footpads, or palpable splenomegaly; or at Day 20 post-infection (the experiment was terminated at Day 16 post-infection as there were no surviving WT or HET mice). Liver-stage infection loads were assessed after injecting them with D-Luciferin in PBS (IVISbrite, PerkinElmer), and imaging using an IVIS Lumina S5 (PerkinElmer).^36^ WT and *Tmprss6*-Het mice were shaved around the abdomen prior to IVIS imaging to account for *Tmprss6*-KO mice having alopecia. Tail blood smears were performed daily from Day 3 post-infection for blood-stage parasitemia assessment. Following euthanasia, blood was collected by cardiac puncture and livers were harvested for RNA extraction and liver iron quantitation.

#### Gene expression analysis

500ng of RNA was reverse transcribed to cDNA using the SensiFAST cDNA synthesis kit (Bioline) following the manufacturer’s protocol. Reverse transcription-qPCR (RT-qPCR) using the SensiFAST SYBR No-ROX kit (Bioline) on a LightCycler 480 II thermocycler (Roche) was used to quantify gene expression levels. Primers and probes used for gene expression analysis and RT-qPCR are listed below:

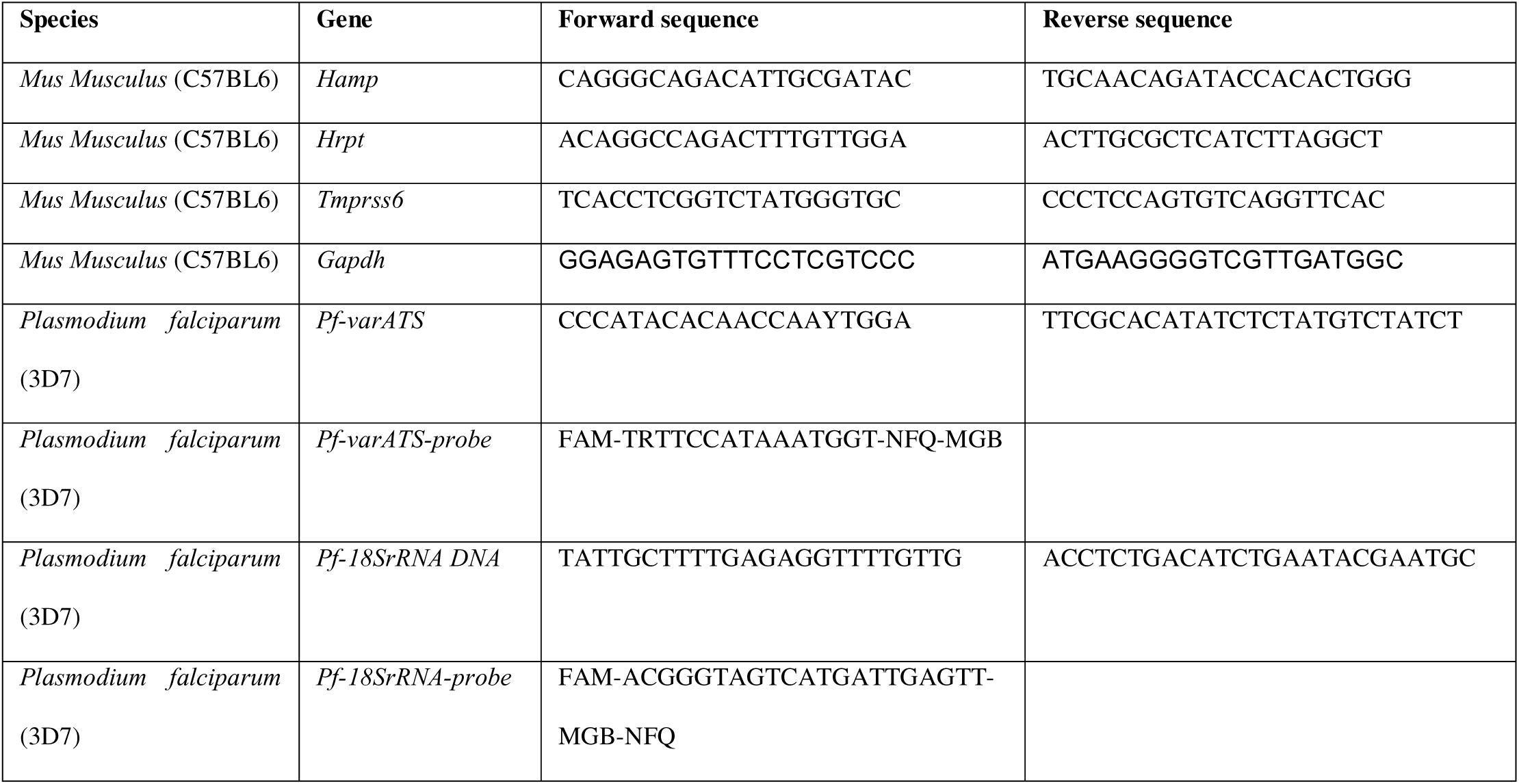

#### Pre-clinical specimen analysis

Liver taken into PBS and then snap frozen at −80°C was used to determine hepatic iron content as previously described.^37^ EDTA anticoagulated murine whole blood was analyzed on an Advia2120i for full blood count determination. Murine plasma iron was determined as previously described.^38^

### In vitro effects of iron restriction on P. falciparum

#### Parasite culture and growth assays

Wild-type *P. falciparum 3D7* asexual blood stage parasites were cultured in Group O RhD+ red blood cells from healthy donors. Growth assays with desferrioxamine (DFO) (Sigma-Aldrich) were set up using sorbitol synchronized ring-stage cultures with a starting parasitemia of ∼1%, assessed by flow cytometry, at 1% hematocrit (HCT) and 50µL volume. At 48 hours parasites were stained with Hoechst and parasitemia analyzed by flow cytometry on a Verse flow cytometer (BD Biosciences, San Jose, USA); Flow cytometry data were analyzed using FlowJo software version 10.0.8 (Tree Star).

Samples for mass spectrometry and RNA sequencing were obtained from parasite cultures synchronized using a combination of serial sorbitol treatments and heparin exposure prior to being set up at early ring-stage (4-6 hours post-invasion) with a ∼1% starting parasitemia, 1% HCT and total volume of 10mL with concentrations of 0 and 12.5µM of DFO. After 30 hours, cultures were treated with saponin to obtain parasite pellets which were frozen at −80°C for proteomic analysis or resuspended in 50µL of RNAlater and stored at −20°C for RNA extraction and sequencing. Four biological replicates were performed at least one week apart, and all samples from the replicates were subsequently processed together.

#### Mass spectrometry

Parasite pellets were lysed in 8 M urea, 100 mM trisaminomethane TRIS (pH 8.8), with protein concentrations determined and normalized by BCA assay (Pierce). Proteins were reduced (10 mM DTT), alkylated (15 mM IAM), and digested overnight with trypsin (1:100). Peptides were acidified, desalted using C18 tips (in house), vacuum-dried, and reconstituted in 2% ACN, 0.1% TFA for LC-MS analysis.

Mass spectrometry was performed on an Orbitrap Fusion Lumos with FAIMS Pro interface (CVs: –40 V and –60 V). MS1 scans were acquired at 120k resolution with AGC target 4 × 10□ (normalized AGC 100%) and max injection time (maxIT) of 50 ms. MS2 scans were acquired at 60k resolution with AGC target 1 × 10□ (normalized AGC 100%) and 50 ms maxIT, All spectra were acquired in positive mode with full scan MS spectra scanning from m/z 300-1600 using a ±1.6 m/z isolation window and CID at 35% collision energy. Peptide separation was performed by nanoflow reversed-phase HPLC (Ultimate 3000 RSLC, Dionex) using an Acclaim PepMap C18 nano-trap column (75 μm × 2 cm) and analytical column (75 μm × 50 cm). The gradient was: 3–23% buffer B over 29 min, 23–40% B in 10 min, 40–80% B in 5 min, held at 80% B for 5 min, then re-equilibrated at 3% B for 10 min. Buffer A: 0.1% FA; buffer B: 80% ACN, 0.1% FA. The mass spectrometry proteomics data have been deposited to the ProteomeXchange Consortium via the PRIDE^39^ partner repository with the dataset identifier PXD064348.

#### RNA sequencing

RNA was extracted using the Isolate II RNA mini kit (Bioline) as per the manufacturer’s protocol. RNA was quantified using the Qubit 2.0 HS RNA assay (Thermo Fisher) and the quality assessed using the 2200 TapeStation System High Sensitivity RNA Screen Tape (Agilent). After normalisation to RNA concentrations of 10ng/µL, sequencing was performed using an in-house mini-bulk sequencing protocol. In short, single-cell transcriptome libraries were generated by adapting the CelsSeq2 protocol^40^ as follows: samples were pooled after first strand cDNA synthesis, treated with Exonuclease 1 for 30 minutes, followed by a 1.2X bead clean-up. Second strand synthesis was performed using NEBNext Second Strand Synthesis module (NEB) in a final reaction volume of 20 µL and NucleoMag NGS Clean-up and Size select magnetic beads (Macherey-Nagel) were used for all DNA purification and size selection steps. Sequencing was performed on the Illumina NextSeq2000 instrument (San Diego, USA), using a 100 cycle kit.

#### Bioinformatic analysis

Briefly, database searching for mass spectrometry data was performed using MaxQuant version 1.6.10.43.^41^ Each raw file was split into two mzXML files using FAIMS-MzXML-Generator for the two CVs used. We assigned the two CVs per raw file as fraction 1 (−40 CV) and fraction 2 (−60CV) across all the files. The *Plasmodium falciparum* 3D7 isolate genome was downloaded from PlasmoDB (release 51)^42^ and used for database searching. For RNA sequencing, expression was quantified by counting the number of unique molecular identifiers (UMIs) and reads mapped to each gene (*P. falciparum* 3D7 genome downloaded from PlasmoDB release 51, and spike-in transcripts (ERCC)) using scPipe (v1.14.0).^43^ Only exonic reads were included in the differential expression analysis.

#### In vitro analyses

Data analysis for mass spectrometry was performed as previously outlined.^44^ Peptide intensities were summed, and non-unique peptides, reverse peptide decoys and contaminants removed. Unique peptides detected in a minimum of 18 samples were log-transformed and quartile-normalised using limma package version 3.46.0 in R,^45^ and missing values imputed using MSImpute using the v1 method.^46^ Differentially expressed peptides between conditions were identified using linear models with empirical Bayes moderated t-statistics. Differentially expressed proteins (DEPs) were estimated using peptide-set enrichment analysis in a gene-set enrichment analysis framework.^47^ For RNA sequencing analysis, differential expression was performed using the negative binomial GLM framework workflow in edgeR (v3.34.0), with enrichment analyses were performed in PlasmoDB. These identified Gene Ontology (GO)^48^ terms and metabolic pathways (from pathway databases - Kyoto Encyclopedia of Genes and Genomes (KEGG)^49^ and MetaCyC^50^.

### Statistical analysis

#### Clinical data

Iron deficiency was defined as ferritin <15µg/L, or <30µg/L if CRP was >5mg/L. Associations between iron status and *P. falciparum* infection and parasitemia at baseline and 28 days post-treatment, as well as the effect of supplementation with IV iron at baseline on the subsequent risk of parasitemia, were analized with Poisson regression models with a log link and robust error variance. Results are presented as point estimates and two-sided 95% confidence intervals (CIs). Additional analyses included adjusted models for pre-specified covariates with the use of directed acyclic graphs as presented in Supplementary figure 1 and 2 and sensitivity analysis excluding women who were malaria RDT positive at screening and were rescreened and deemed malaria microscopy-slide negative at enrolment. For analysis of parasite densities across timepoints by treatment group, we calculated median values with interquartile ranges (IQRs). Due to the skewed nature of the data, parasite densities were log-transformed and groups were compared using appropriate statistical testing. Analysis were performed using Stata SE, version 18.0 (StataCorp, College Station, TX).

#### Pre-clinical models

Sample sizes for animal experiments were calculated based on expected differences in survival between groups with 80% power at an alpha of 0.05, accounting for expected effect sizes based on preliminary data. Statistical testing of continuous variables was performed using Ordinary One-way ANOVA with Dunnet correction for multiple comparisons (except Figure 2e, where a Kruskal-Wallis test with Dunn’s correction for multiple comparisons was used) or Two-way Repeated Measures ANOVA with Dunnett’s correction for multiple comparisons. Survival analyses were performed using Gehan-Breslow-Wilcoxon tests. Statistical analyses were performed in Prism version 10.4.1 (GraphPad software).

**Figure 2.**
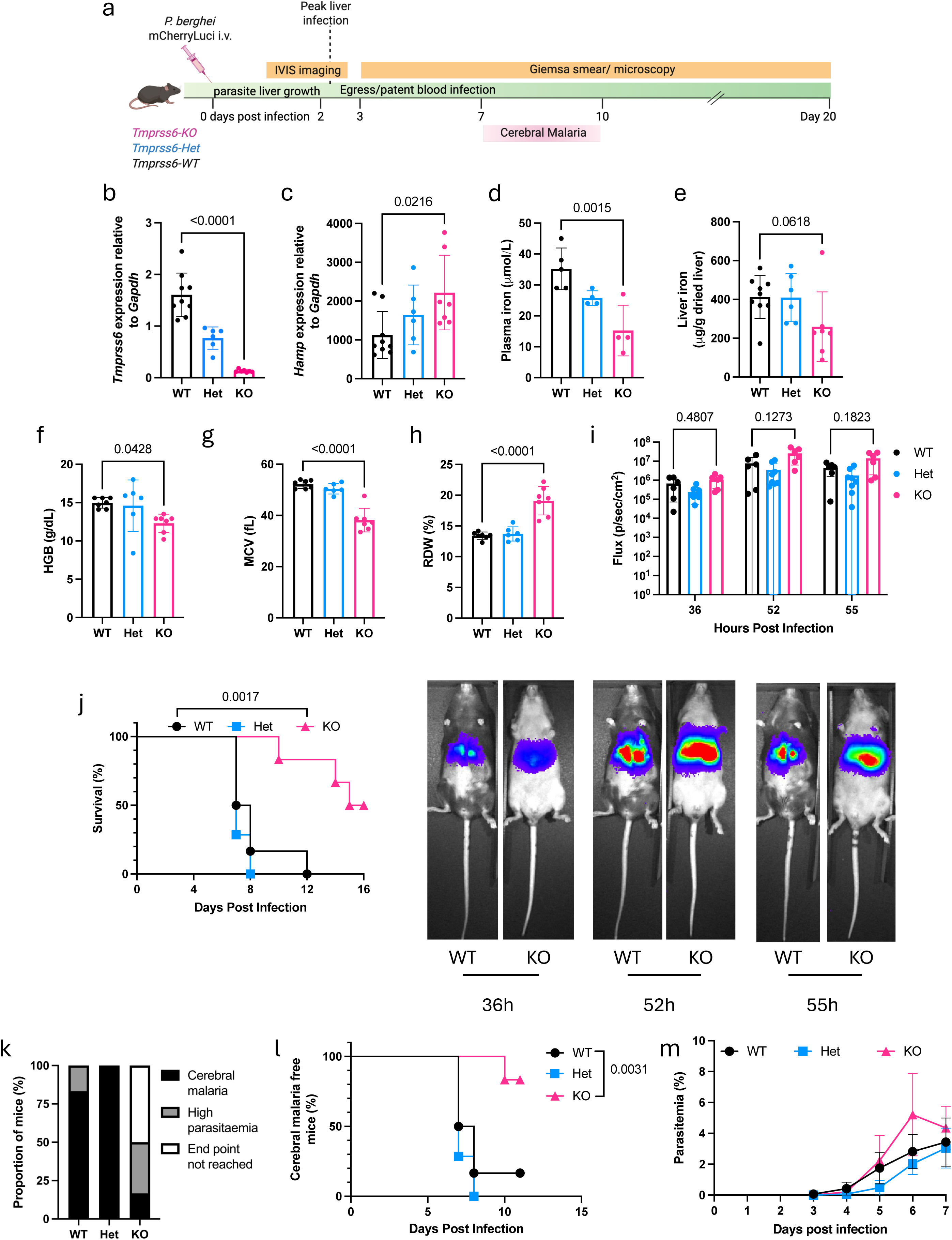
Iron deficiency in *Tmprss6*-knockout mice provides protection against *P. berghei* infection. (a) Experimental design for *P. berghei* infection in C57BL/6 wild-type (WT), *Tmprss6*-heterozygous (Het), and *Tmprss6*-knockout (KO) mice. Mice reach cerebral malaria within 7-10 days post-infection. Created with Biorender.com (b) Hepatic *Tmprss6* mRNA expression in uninfected mice. (c) Hepatic hepcidin (*Hamp*) mRNA expression. (d) Plasma iron concentration. (e) Non-heme liver iron content. (f-h) Hematological parameters: (f) hemoglobin concentration (HGB), (g) mean corpuscular volume (MCV), and (h) red cell distribution width (RDW). (i) IVIS imaging of liver-stage infection showing parasite burden in all mice at indicated hours post-infection. Representative IVIS images of select mice are shown below. (j) Kaplan-Meier survival curves following *P. berghei* infection. (k) Percentage of mice succumbing to cerebral malaria (black), high parasitemia (grey) or not reaching humane endpoint (white). (l) Kaplan-Meier survival curve of mice succumbing to cerebral malaria. (m) Blood-stage parasitemia. Data represented as mean ± SD, n=6-9 mice per group (b-c and e), n=4-5 (d), n=6-7 (f-h and j-m), and n=6-8 (I).

## Results

### Sample characteristics

We evaluated a total of 3,141 samples, collected from 711 women in Zomba, Southern Malawi, across 5 timepoints during the trial (Figure 1a, Supplementary Table 1). All enrolled women were anemic by capillary hemoglobin (Hb), with Hb<10 g/dL. Cohort characteristics were similar between time points, despite not all participants having samples at all timepoints (Supplementary Table 1). At enrolment, 55.9% (358/641) of women with available samples were primigravid and there was a relatively high percentage of women with HIV infection (14.2%; 90/634), which was expected in this setting. Iron-deficiency and inflammation were common, with 39.1% (245/626) of women being iron-deficient and 52.7% (330/626) being inflamed.

### Plasmodium falciparum infection by malaria RDT at enrolment

One third (33.1%, 212/641) of the women included in this study were malaria positive at pre-screening by RDT performed on a finger-prick sample. These women were treated with artemether-lumefantrine (AL), rescreened (no sooner than one week after AL treatment) and malaria-free by malaria microscopy slide (Figure 1a, Supplementary Table 1). Once enrolled, all women had a venous sample taken, which was subsequently used to repeat the malaria RDT—1.4% (9/627) of samples had a venous sample positive RDT result.

### Plasmodium falciparum infection by qPCR

We assessed the prevalence of *P. falciparum* on venous whole blood at all time points by ultra-sensitive qPCR (Figure 1b). A total of 284/641 (44.3%) of samples were *P. falciparum* positive by qPCR at enrolment. Excluding from this analysis all the women who were malaria RDT positive at screening and allowed to enter the trial after being rescreened (after AL treatment and with possible dead parasites or parasite DNA still in circulation) that prevalence dropped to 35.9% (154/429) (Figure 1b). As all women who were not rescreened entered the trial after a negative malaria RDT, this shows that a high-proportion of *P. falciparum* infections go undetected by RDT at this time point in pregnancy in this area of Malawi. After enrolment, and receipt of the anti-malarial sulfadoxine-pyrimethamine (IPTp-SP), the overall prevalence of positive qPCR for *P. falciparum* dropped to 18.2% (120/659) at 28 days post-treatment and 11.9% (46/386) at delivery (Figure 1b). The prevalence was similar between those who entered the trial directly or after rescreen. The overall prevalence of *P. falciparum* positive qPCR remained at 10.8% (32/295) after delivery and into the 28 days postpartum time point. It is worth noting that 81/284 (28.5%) of women with a positive qPCR result at baseline were also qPCR positive 28 days post-treatment. We were not able to assess if these were persisting infections or new infections. At baseline, parasite densities in those who were qPCR positive, were expectedly low (median (IQR) 1.01 (0.22-9.48) parasites/μL) and remained so throughout all time points (Figure 1c). Of those enrolled with a negative RDT (not rescreened), 11.6% (20/171) were positive by qPCR, possibly representing RDT false-negatives (Supplementary Figure 3 – blue dots). Those who entered the study with a microscopy-negative slide after rescreening but with a positive qPCR, may have had true sub-microscopic infections, or likely had circulating dead parasites or parasite DNA (Supplementary Figure 3 – red dots).

### Impact of iron status on prevalence and density of parasitemia

We sought to understand the associations between iron status and *P. falciparum* infection and parasitemia. In the REVAMP women, at baseline, iron-deficiency was associated with a 63% reduction in the probability of being *P. falciparum* qPCR positive (95% CI [51%-73%]) (Table 1 and Supplementary table 2). This association remained strong when adjusted for gestational age at baseline, gravidity status, HIV, rescreened + AL treatment, maternal age, education, income source, and religion (50% reduced risk, [30%-64%]) (Table 1). This association was maintained when excluding the women who were rescreened (55% adjusted reduced prevalence, 95% CI [34%-69%]) (Table 1 and Supplementary table 3).

**Table 1.**
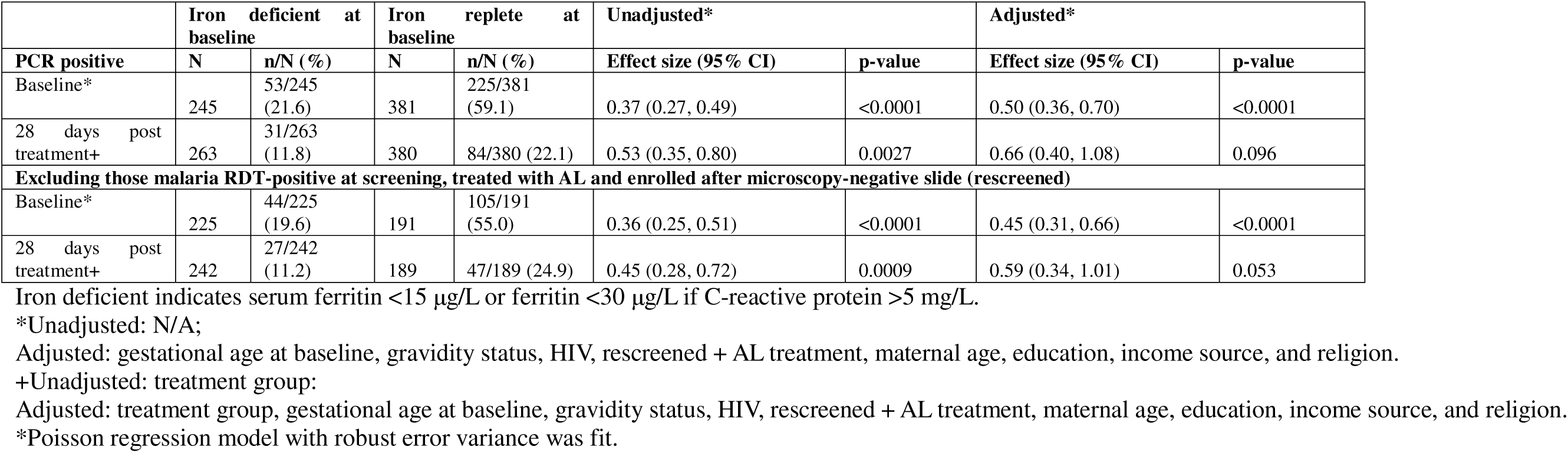
Comparison by ID/IR at baseline and qPCR positivity.

|  | Iron deficient at baseline |  | Iron replete at baseline |  | Unadjusted* |  | Adjusted* |  |
| --- | --- | --- | --- | --- | --- | --- | --- | --- |
|  | N | n/N (%) | N | n/N (%) | Effect size (95% CI) | p-value | Effect size (95% CI) | p-value |
| **PCR positive** |  |  |  |  |  |  |  |  |
| Baseline* | 245 | 53/245 (21.6) | 381 | 225/381 (59.1) | 0.37 (0.27, 0.49) | <0.0001 | 0.50 (0.36, 0.70) | <0.0001 |
| 28 days post treatment+ | 263 | 31/263 (11.8) | 380 | 84/380 (22.1) | 0.53 (0.35, 0.80) | 0.0027 | 0.66 (0.40, 1.08) | 0.096 |
| **Excluding those malaria RDT-positive at screening, treated with AL and enrolled after microscopy-negative slide (rescreened)** |  |  |  |  |  |  |  |  |
| Baseline* | 225 | 44/225 (19.6) | 191 | 105/191 (55.0) | 0.36 (0.25, 0.51) | <0.0001 | 0.45 (0.31, 0.66) | <0.0001 |
| 28 days post treatment+ | 242 | 27/242 (11.2) | 189 | 47/189 (24.9) | 0.45 (0.28, 0.72) | 0.0009 | 0.59 (0.34, 1.01) | 0.053 |

Iron deficient indicates serum ferritin <15 µg/L or ferritin <30 µg/L if C-reactive protein >5 mg/L.
\*Unadjusted: N/A;
Adjusted: gestational age at baseline, gravidity status, HIV, rescreened + AL treatment, maternal age, education, income source, and religion.
+Unadjusted: treatment group:
Adjusted: treatment group, gestational age at baseline, gravidity status, HIV, rescreened + AL treatment, maternal age, education, income source, and religion.
\*Poisson regression model with robust error variance was fit.

We looked at the association of *P. falciparum* qPCR positivity at 28 days post treatment, considering iron-deficiency status at baseline (Table 1). The strength and direction of these associations were maintained after adjustment but did not reach statistical significance (prevalence reductions of 34% [−0.8%, 60%], and 41% [−0.1%, 66%], respectively) (Table 1). Sensitivity analysis, performed to include the WHO definition of iron deficiency (serum ferritin <15 μg/L or ferritin <70 μg/L if CRP>5 mg/L), revealed similar patterns (Supplementary Table 2 and 4). We found similar strength and direction of associations when compared to the main analysis. Additionally, we conducted sensitivity analysis of the association between qPCR positivity at 28 days post treatment and iron status at baseline in only those women entering the trial with a negative qPCR result at baseline, thus assessing the risk of becoming infected (Supplementary table 5). We found no evidence of an effect of iron status at baseline on the risk of becoming qPCR positive (risk reductions ID versus IR: 5% [−119%, 59%], p=0.91).

Overall quantification of parasitemia in those who were qPCR positive, revealed similar parasite densities irrespective of iron status at both enrolment (mean difference [95%CI] −0.16 [−0.91, 0.60], p = 0.69) and 28 days post treatment (−0.17 [−2.75, 0.40], p=0.14) (Figure 1d). Similar analysis of parasitemia on those individuals not rescreened or in those coming into the trial after being rescreened revealed baseline iron-status was not associated with parasite densities at baseline or at the 28 days post-treatment visit (Figure 1 e-f). Sensitivity analysis of the parasite densities at 28 days post-treatment by iron status at baseline in those women with a qPCR negative result at baseline revealed a higher median parasite density in iron replete versus iron deficient women (19.60 (0.33-558.50) parasites/μL versus 3.27 (0.70-439.66) parasites/μL) (Supplementary table 6).

### Impact of iron intervention on prevalence and density of parasitemia

We next investigated the effect of supplementation with IV iron at baseline compared to oral iron on the subsequent prevalence of parasitemia (Table 2). Treatment with IV Iron was not associated with increased prevalence of parasitemia positivity by qPCR at any of the subsequent visits when compared to oral iron. Adjusting the analysis for iron-status at previous visit and time between visits (among others) did not alter the risk of becoming qPCR positive. The same pattern was observed when excluding from the analysis those individuals who were enrolled after rescreening (Table 2).

**Table 2.** Comparison of qPCR positivity by treatment group (IV iron versus Oral iron)

| PCR positive | IV Iron |  | Oral Iron |  | Unadjusted * |  | Adjusted* |  |
| --- | --- | --- | --- | --- | --- | --- | --- | --- |
|  | N | n/N (%) | N | n/N (%) | Effect size (95% CI) | p-value | Effect size (95% CI) | p-value |
| 28 days post treatment | 334 | 65/334 (19.5) | 325 | 55/325 (16.9) | 1.15 (0.83, 1.59) | 0.40 | 1.14 (0.81, 1.61) | 0.46 |
| 36 weeks' gestation | 182 | 26/182 (14.3) | 195 | 23/195 (11.8) | 1.21 (0.72, 2.05) | 0.47 | 1.45 (0.77, 2.74) | 0.25 |
| Delivery | 191 | 17/191 (8.9) | 195 | 29/195 (14.9) | 0.60 (0.34, 1.05) | 0.072 | 0.56 (0.30, 1.02) | 0.059 |
| 28 days postpartum | 138 | 10/138 (7.2) | 157 | 22/157 (14.0) | 0.52 (0.25, 1.05) | 0.070 | 0.52 (0.24, 1.14)+ | 0.10 |
| **Excluding those malaria RDT-positive at screening, treated with AL and enrolled after microscopy-negative slide (rescreened)** |  |  |  |  |  |  |  |  |
| 28 days post treatment | 226 | 40/226 (17.7) | 219 | 38/219 (17.4) | 1.02 (0.68, 1.53) | 0.92 | 0.95 (0.62, 1.44) | 0.81 |
| 36 weeks' gestation | 117 | 13/117 (11.1) | 131 | 14/131 (10.7) | 1.04 (0.51, 2.12) | 0.91 | 1.86 (0.65, 5.31) | 0.25 |
| Delivery | 120 | 10/120 (8.3) | 131 | 17/131 (13.0) | 0.64 (0.31, 1.35) | 0.24 | 0.67 (0.31, 1.45) | 0.31 |
| 28 days postpartum | 85 | 5/85 (5.9) | 106 | 16/106 (15.1) | 0.39 (0.15, 1.02) | 0.056 | 0.38 (0.14, 1.08)+ | 0.070 |

Adjusted: ID at previous visit, anemia at previous visit, HIV status, rescreened + AL treatment, gravidity, maternal age, concurrent gestational age, time between visits.
+: As “Adjusted”, excluding concurrent gestational age.
\*Poisson regression model with robust error variance was fit (by timepoint). The model estimates the direct effect, not total effect given ID at previous visit, and anemia at previous visit are mediators in the model.

An analysis of the parasite densities across all visits by treatment group revealed higher parasite densities in the IV Iron group at 28 days post-treatment (N, median (IQR) N=86, 5.66 (0.12 – 601.13)) compared to the oral iron group (N=85, 0.36 (0.08 – 33.23) (Figure 1g). However, this pattern was inverted at delivery and 28 days postpartum, with individuals in the IV Iron group presenting reduced parasite densities (Figure 1g).

### Genetic dissection of iron homeostasis in pre-clinical models of malaria

Interpreting the effects of iron status on *Plasmodium* risk in a field cohort is complex. Animal models of *Plasmodium* infection allow us to better dissect the timing and effect of iron status. To isolate the effects of iron status on *Plasmodium* infection and disease burden, we used *P. berghei* infection to induce liver and blood-stage infection, as well as cerebral malaria, in a mouse model of iron deficiency (*Tmprss6*-KO mice) (Figure 2a).

Inactivation of *Tmprss6* causes an inability to suppress hepcidin expression, leading to reduced iron absorption and utilization, plasma iron depletion and iron deficiency phenotype.^51^ Uninfected *Tmprss6*-KO mice had ablated hepatic *Tmprss6* mRNA expression (Figure 2b), leading to increased hepcidin expression (Figure 2c). *Tmprss6*-KO mice showed reduced plasma iron (Figure 2d), and a non-statistically significant reduction in hepatic iron (Figure 2e). As expected, *Tmprss6*-KO mice displayed iron restricted erythropoiesis with decreased hemoglobin (HGB) concentrations and mean corpuscular volumes (MCVs) and increased red cell distribution width (RDW) compared to WT mice (Figure 2f-h). *Tmprss6*-Het mice have hepatic *Tmprss6* expression mid-way between WT and *Tmprss6*-KO animals with a complementary increase in hepcidin (*Hamp*) mRNA mid-way between WT and *Tmprss6*-KO (Figure 2b-c). However, *Tmprss6*-Het mice display iron and erythroid parameters almost identical to WT animals.

IVIS imaging of animals infected with *P. berghei* sporozoites expressing mCherry and luciferase revealed *Tmprss6*-KO mice had comparable intensity of liver infection to that of both WT and *Tmprss6*-Het mice from 36 to 55 hours post-infection (Figure 2i). In this model of *P. berghei* infection, animals display cerebral malaria symptoms between days 7 and 10 post-infection. *Tmprss6*-KO mice experienced significantly improved survival compared to *Tmprss6*-Het and WT animals. The median survival for *P. berghei* infected *Tmprss6*-KO mice was 15.5 days compared with 7.5 days and 7 days for *Tmprss6*-Het and WT mice, respectively (p<0.002) (Figure 2j). Survival from cerebral malaria was also significantly improved in *Tmprss6*-KO mice (83% survival in *Tmprss6*-KO animals vs 17% WT; Figure 2k) and time to succumb to cerebral malaria was delayed (median time to cerebral malaria: 10 days *Tmprss6*-KO vs 7 days WT; p<0.005 Figure 2l). However, parasitemia from day 3 to day 7 (the timepoint at which most WT and *Tmprss6*-Het mice succumbed to cerebral malaria), was similar between *Tmprss6*-KO and WT animals (Figure 2m). Parasitemia was higher in the only WT mouse that survived cerebral malaria, when compared with the 5 surviving *Tmprss6*-KO mice (Supplementary figure 4).

### Direct effects of iron deprivation on Plasmodium falciparum in vitro

Having observed a protective effect of host iron deficiency on *Plasmodium* infection in both our clinical and pre-clinical models, we finally sought to establish whether iron chelation in parasite cultures functionally influences *Plasmodium* molecular profiles.

The antimalarial actions of iron chelators on blood-stage *Plasmodium* parasites have been previously identified but the full molecular effects of this chelation have not been thoroughly defined.^52^ We used the iron chelator desferrioxamine (DFO) in 3D7 *P. falciparum* blood-stage cultures followed by transcriptomic and proteomic analysis to investigate the effect of iron restriction (Figure 3a). To confirm an antimalarial action of DFO, we first exposed *in vitro* cultures of 3D7 *P. falciparum* to DFO for 48 hours; a dose dependent inhibition of parasite growth was evident (Figure 3b). Similar to previous reports, we found the IC50 for DFO to be 16.16µM (95% CI 14.99 – 17.33).^53,54^

**Figure 3.**
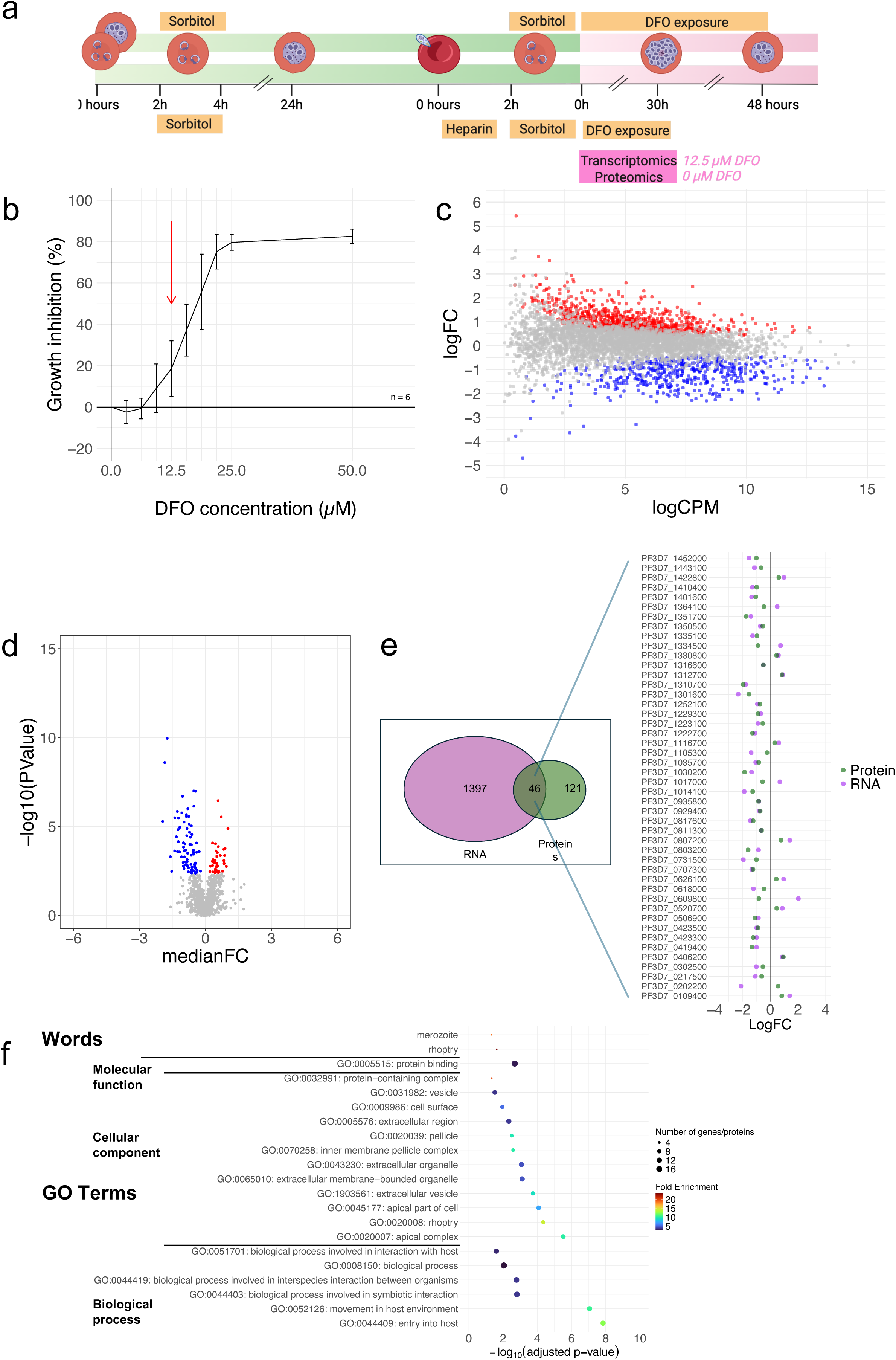
Iron chelation slows parasite development and induces extensive transcriptomic and proteomic changes in *P. falciparum*. (a) Experimental design for in vitro analysis of iron chelation on *P. berghei.* Created with Biorender.com (b) Dose-dependent growth inhibition of 3D7 *P. falciparum* parasites exposed to the iron chelator desferrioxamine (DFO) for 48 hours. (c) Volcano plot showing differentially expressed genes in parasites treated with 12.5μM DFO for 30 hours (889 upregulated [red], 508 downregulated [blue] genes; log□ fold change ≥1, false discovery rate <0.05). (d) Volcano plot of differentially expressed proteins (40 upregulated, 81 downregulated; log□ fold change ≥1, false discovery rate <0.05). (e) Overlap between differentially expressed genes and proteins, showing concordant and discordant regulation. (f) Enriched Gene Ontology (GO) biological process terms for the 46 differentially expressed gene-protein pairs, highlighting processes involved in host cell interaction and invasion. (b) Data represented as mean ± SD, n=6.

To characterise the effect of iron chelation on the transcriptome and proteome of the parasite, we performed RNA sequencing and quantitative proteomics on synchronous WT 3D7 *P. falciparum* cultures exposed to 12.5µM of DFO for 30 hours from early ring-stage (4-6 hours post invasion, Figure 3a), a concentration of DFO expected to affect parasite growth, without causing excessive parasite death that could render results difficult to interpret.

A total of 1,397 genes were significantly differentially expressed (FDR<0.05) in cultures exposed to DFO for 30 hours compared with control parasites in media alone, of which 889 were upregulated and 508 downregulated (Figure 3c). At the protein level, 121 proteins were differentially expressed in DFO treated parasites compared with controls comprising 81 proteins significantly downregulated and 40 upregulated (Figure 3d). Of these, 46 were differentially expressed at both the RNA and protein level. Most commonly this differential expression was in the same direction – 32 downregulated in both, 9 upregulated in both (Figure 3e). Among the 46 gene/protein pairs differentially expressed in both analyses, 15 are associated with merozoite invasion,^55^ 5 are characterized as exported proteins,^56^ 6 are associated with the inner membrane complex,^57,58^ and 4 are associated with the food vacuole.^59^ Enrichment analysis performed for genes/proteins differentially expressed at both an RNA and protein level revealed enriched biological process GO terms associated with interaction with the host and entry into host (Figure 3f).

## Discussion

Our multilayered approach to understanding the relationship between iron-deficiency and *P. falciparum* infection has yielded consistent findings across clinical, pre-clinical and *in vitro* data. In Malawian pregnant women in the REVAMP trial—an RCT of IV iron versus standard of care oral iron in anemic women—, iron deficiency at baseline in the second trimester, and prior to iron intervention, was associated with a 50% reduced probability of being qPCR positive for *P. falciparum*. Additionally, iron-deficiency at baseline was associated with a lower probability of *P. falciparum* parasitemia at 28 days post enrolment. These associations were maintained when looking exclusively at women who were malaria RDT-negative at pre-screening, suggesting an effect that is independent from recent malaria treatment. Treatment with IV iron was not associated with an increased prevalence of parasitemia at any timepoint during the trial when compared with the standard of care, oral iron treatment. Complementing these clinical observations, our pre-clinical models using iron-deficient *Tmprss6-*knockout mice exhibited improved survival and reduced cerebral malaria following *P. berghei* infection. Finally, *in vitro* cultures using 3D7 parasites confirmed these antimalarial effects at the molecular level with iron chelation resulting in substantial inhibition of parasite growth and substantial changes in the transcriptome and proteome of *P. falciparum*. Together, our data all converges to support the protective role of iron deficiency against *Plasmodium* infection and disease progression.

Previous studies in pregnant women have been contradictory about the protective effects of iron deficiency.^60,61^ Some of that contradiction may be derived from the definition of iron deficiency used in any particular study, as well as from the underlying characteristics of the women; however, a protective association of iron deficiency against *Plasmodium* infection is consistently observed when using ferritin as a biomarker of iron stores.^61^ The strong protective association we observed for maternal peripheral infections in the second trimester (50% reduced risk, 95% CI [30%-64%], p<0.0001) is of similar magnitude to the one found in a case-control study conducted in a similar area of Malawi in which placental malaria infections (measured at delivery) were found to be less frequent in women who were iron-deficient (OR 0.4 [0.2-0.8], p = 0.01).^62^ Also, a secondary analysis of an antimalarial RCT conducted in Papua New Guinea—where the mean hemoglobin at enrolment during the second trimester was 9.7 g/dL— found women who were iron deficient at enrolment were 50% less likely to be *Plasmodium* positive by microscopy (adjusted odds ratio [aOR] 0.50 [0.38, 0.66], P□<□0.001).^28^ While most studies make concurrent observations of iron deficiency and *Plasmodium* infection—or look at association between time points that are extremely far apart (second trimester versus delivery)—and are thus unable to determine the direction of the association, we determined the influence of iron deficiency at baseline with risk of *Plasmodium* infection 28 days after. Although all women in our cohort received iron treatment (either IV or oral) the iron utilization over four weeks would still be different between iron-deficient and replete women. In women overall, as well as excluding those women who received AL treatment before entering the study, iron deficiency at baseline was still more likely to protect from *Plasmodium* infection 28 days after (Table 1). Interestingly, among women who were qPCR negative at baseline, iron status did not influence the risk of qPCR positivity at 28 days post treatment, likely reflecting the stronger effect of being on trial (receipt of insecticide treated nets, and antimalarial drugs) compared to the effect of iron status (Supplementary table 3). Importantly, while iron deficiency was protective overall, our analysis of IV iron versus oral iron supplementation showed no significant difference in subsequent parasitemia risk by qPCR across treatment groups. This mirrors the findings of the main REVAMP trial^29^ (which evaluated clinical malaria cases) and those of other iron-supplementation RCTs in malaria endemic areas^24^ and suggests that when appropriate malaria prevention measures are in place (all women received IPTp-SP or cotrimoxazole), iron supplementation— independently of the route of administration—can be given safely, supporting current WHO recommendations for universal iron supplementation in high-anemia-prevalence settings.^63^ However, the transiently higher parasite densities observed in the IV iron group at 28 days post-treatment highlight the need for vigilant malaria surveillance in the period immediately following iron repletion.

Our mouse model of iron deficiency — genetic knockout of *Tmprss6, a* gene regulating systemic iron homeostasis^33,51^—provides important mechanistic insights that are difficult to disentangle in human populations. Additionally, our pre-clinical model of infection with sporozoites, replicates human infection—unlike previous studies that have bypassed liver-stage infection using infected red cells.^64–68^ After infection with *P. berghei* sporozoites, *Tmprss6*-knockout mice displayed comparable liver infection intensity (Figure 2i) despite reduced hepatic iron content (Figure 2e), suggesting iron may not be critical for *Plasmodium* growth in the liver, or that minimal liver iron requirements for *Plasmodium* growth are below the reduction achieved in the model. Others have observed a protective effect of acute elevated hepcidin coupled with prior inflammation on liver infection burden caused by *P. berghei* sporozoites,^69^ however the chronic elevated hepcidin in the absence of prior inflammation seen in our model did not exhibit the same liver-protection effect. Following *Plasmodium* egress from the liver, *Tmprss6*-KO animals displayed improved survival and protection from cerebral malaria. It has been shown that *in vitro* culture of 3D7 parasites in erythrocytes from iron-deficient donors result in lower parasitaemia when compared to 3D7 grown in erythrocytes from iron-replete donors.^70^ The early development of blood-stage infection in *Tmprss6*-KO mice, in conditions where iron availability is restricted and erythropoiesis shows an iron-deficient phenotype (Figure 2d,f-h), did not differ (Figure 2m) from WT mice. This mirrors the similar parasite densities observed in women who were iron deficient compared to those who were iron replete in our clinical cohort (Figure 1d). The low numbers of WT mice surviving past the cerebral malaria window in this model of infection (N=1) do not allow us to make conclusions regarding a possible parasitemia control in *Tmprss6*-KO mice past day 7 (Supplementary Figure 4). We observed a significant increase in survival (Figure 2j-k) and a delayed time to cerebral malaria in *Tmprss6*-KO mice (median 10 days versus 7 days in WT mice, p<0.005) suggesting that iron restriction leading to iron-restricted erythropoiesis impacts the pathogenic processes leading to severe disease manifestations (a finding that can have important clinical implications, as previously reported).^71^ The mechanisms of action of iron-restriction on clinical symptoms could involve altered cytoadherence properties, reduced inflammatory responses, or changes in endothelial activation—all critical factors in cerebral malaria pathogenesis.^72,73^ This lack of effect on parasitemia in our mouse experiments mirrors what we observed in our clinical cohort, where iron-deficiency was associated with less risk of infection, but not with parasite density.

Iron chelation with DFO is not only an *in vitro* tool but also a clinically relevant one.^71^ Our transcriptomic and proteomic analyses of *P. falciparum* cultured under iron-chelated conditions reveal the molecular adaptations that may underlie the parasite’s response to systemic iron restriction. Our transcriptome and proteome wide analysis of DFO-exposed *P. falciparum* parasite cultures identified substantial changes in mRNA and protein expression by iron chelation. We identified 1,397 differentially expressed genes and 121 differentially expressed proteins in DFO-treated parasites, with 46 entities showing mostly concordant changes at both RNA and protein levels (41/46 pairs). Similar to the transcriptomic findings of 3D7 grown in iron deficient erythrocytes,^70^ we identified changes in proteins and genes involved in merozoite invasion, protein export and RNA binding, and associated with the food vacuole – processes and organelles important for parasite survival, virulence and nutrient acquisition. The enrichment of GO terms related to host interaction and cell entry further supports the hypothesis that iron deficiency primarily protects by impairing the parasite’s ability to invade and establish infection within host cells. This molecular *in vitro* evidence bridges our clinical observation of reduced infection prevalence in iron-deficient women with the improved survival seen in iron-deficient mice. These data highlight iron as an important nutrient for *P. falciparum* and indicate proteins of interest for future study.

While our manuscript focussed primarily on the effects of iron on *Plasmodium* biology, there is an element of interaction between iron and immunity that represents another interface that can affect the relationship between iron-deficiency and infection. Outside pregnancy, iron’s essential role in immune function is well-established ^74,75^. Recent evidence shows that iron status influences antibody responses to vaccines ^76^ and may affect antibody functionality. ^77^ In pregnancy, both the inflammatory and antibody response to *Plasmodium* are key elements that determine the course of the infection. ^21,78^ In our pregnant clinical cohort, pre-existing immunity (or lack of) from previous malaria exposures may interact with iron deficiency status. The reduced parasitemia risk we observed in iron-deficient women might reflect an optimal balance where modest reductions in adaptive immunity are offset by direct inhibitory effects on parasite growth and virulence. Additionally, our transcriptomic and proteomic analyses under iron-restricted conditions revealed significant changes in parasite-exported proteins, as well as proteins essential for erythrocyte invasion and intraerythrocytic development (Supplementary table 3). Many of these proteins are known to interact with the host immune system and modify infected erythrocyte surfaces, thereby affecting immunogenicity and cytoadherence properties. ^79,80^ While these potential immune-related mechanisms provide additional context for our findings, they remain largely speculative and highlight important areas for future research.

Our findings have several important implications for public health policy and clinical practice. Our data provide further evidence for the protective effect of iron deficiency on *Plasmodium* infection. The absence of increased infection risk in our IV iron treatment group (compared to standard of care oral iron) suggests that rapidly correcting iron deficiency can be done safely in anemic women when appropriate malaria control measures are in place. However, this reinforces the necessity of careful consideration when implementing iron supplementation programs in malaria endemic areas. While correcting anemia remains a critical goal, particularly during pregnancy, iron repletion may transiently increase susceptibility to malaria highlighting the need for enhanced vigilance and possibly intensified malaria prevention during the period immediately following iron administration. Finally, our pre-clinical and molecular findings suggest that under iron-restriction conditions there are substantial transcriptomic and proteomic changes that are essential for parasite survival and pathogenicity. These can potentially be targeted as new therapeutics avenues.

Our study has several limitations that should be considered. In our clinical cohort, all women enrolled were anemic and received one form of iron supplementation. While our models were adjusted for measured well-known confounders of the associations, estimates may be biased by the presence of unmeasured confounders. Additionally, the numbers of women with longitudinal data available across the all trial did not allow for the necessary statistical power to model the interaction between iron-deficiency and *Plasmodium* infection throughout. We also did not have enough infected women to evaluate the influence of new infections versus persistent infections in this cohort. In our pre-clinical model, while the *Tmprss6*-knockout approach offers several advantages for isolating the impact of iron deficiency, it does not perfectly recapitulate the complex and often multifactorial iron deficiency observed in pregnant human populations. Future studies could explore a range of iron deficiency models, including pregnancy models, with varying severity and mechanisms to better understand dose-response relationships. Our in vitro experiments used DFO as an iron chelator, which may have effects beyond just iron restriction and may not replicate the iron-restricted erythropoiesis that is characteristic of iron deficiency. Alternative approaches using direct manipulation of parasite iron-related genes would strengthen the evidence regarding the role of iron in parasite biology.

In conclusion, our study demonstrates that iron deficiency protects against *Plasmodium* infection across clinical, pre-clinical, and molecular contexts. In anemic pregnant Malawian women, iron deficiency reduced *P. falciparum* parasitemia risk, while iron-deficient mice showed improved survival against cerebral malaria. Molecular analyses of DFO-exposed parasite cultures revealed iron restriction significantly alters parasite transcriptomes and proteomes, particularly affecting invasion pathways. Future research should explore iron-dependent parasite metabolic pathways as potential therapeutic targets and optimise iron supplementation protocols to balance maternal health benefits with malaria risk in vulnerable populations.

## Acknowledgments

The REVAMP trial was funded by the Bill and Melinda Gates Foundation (INV-010612). S.-R.P. is funded by a National Health and Medical Research Council of Australia (NHMRC) Fellowship (GNT1158696 and GNT2009047). C.B. is funded by the NHMRC Fellowship (GNT2033705). JAB was funded by the NHMRC (1139153, 1176955) and is an NHMRC Leadership Fellow. W-H.T. is supported by an NHMRC Investigator Grant GNT2016908, Victorian State Government Operational Infrastructure Support and an NHMRC Independent Research Institutes Infrastructure Support Scheme (IRIISS). We thank Malawian health workers, participants and their families for their involvement in the REVAMP study. We thank the WEHI animal house staff and the WEHI Insectary facility.

## Data availability

The mass spectrometry proteomics data have been deposited to the ProteomeXchange Consortium via the PRIDE^39^ partner repository with the dataset identifier PXD064348. Underlying de-identified individual participant data encompassing the clinical data results and a data dictionary are accessible at figshare (*data available from the corresponding author upon request).* All in vitro transcriptomic RNA-Seq data relevant to the paper are available on Gene Expression Omnibus (*data available from the corresponding author upon request*).

