## Supplementary data for "Host iron deficiency protects against *Plasmodium* infection and drives parasite molecular reprofiling"

**Supplementary Table 1. Baseline Characteristics for those with non-missing parasitemia across all visits**

| | Baseline | 28 days post treatment | 36 weeks gestation | Delivery | 28 days postpartum |
|---|---|---|---|---|---|
| | N=641 | N=659 | N=377 | N=385 | N=295 |
| Age (years) | 22·0 (6·1) | 22·0 (6·1) | 22·6 (6·3) | 22·7 (6·4) | 22·9 (6·5) |
| Primigravid | 358/641 (55·9%) | 370/659 (56·1%) | 196/377 (52·0%) | 201/385 (52·2%) | 148/295 (50·2%) |
| Gestational age (weeks) | 22·0 (19·6-24·3) | 22·0 (19·6-24·1) | 22·4 (19·7-24·4) | 22·1 (19·6-24·6) | 22·4 (19·9-24·6) |
| Height (cm) | 155·3 (6·5) | 155·2 (6·5) | 155·7 (6·7) | 155·8 (6·4) | 155·9 (6·9) |
| Weight (kg) | 55·3 (8·2) | 55·3 (8·2) | 55·6 (8·3) | 55·7 (8·1) | 56·0 (8·1) |
| Body Mass Index (kg/m2) † | 22·9 (3·0) | 22·9 (3·0) | 22·9 (3·0) | 22·9 (2·9) | 23·0 (2·9) |
| Religion ‡ | | | | | |
| None | 2/638 (0·3%) | 2/656 (0·3%) | 0/375 (0·0%) | 0/383 (0·0%) | 0/293 (0·0%) |
| Christian | 438/638 (68·7%) | 449/656 (68·4%) | 256/375 (68·3%) | 266/383 (69·5%) | 201/293 (68·6%) |
| Muslim | 193/638 (30·3%) | 200/656 (30·5%) | 118/375 (31·5%) | 115/383 (30·0%) | 91/293 (31·1%) |
| Other | 5/638 (0·8%) | 5/656 (0·8%) | 1/375 (0·3%) | 2/383 (0·5%) | 1/293 (0·3%) |
| Education ‡ | | | | | |
| None | 2/620 (0·3%) | 2/636 (0·3%) | 1/364 (0·3%) | 2/373 (0·5%) | 1/284 (0·4%) |
| Lower Primary(1-5) | 146/620 (23·5%) | 146/636 (23·0%) | 75/364 (20·6%) | 73/373 (19·6%) | 53/284 (18·7%) |
| Upper Primary(6-8) | 269/620 (43·4%) | 279/636 (43·9%) | 153/364 (42·0%) | 164/373 (44·0%) | 119/284 (41·9%) |
| Lower Secondary(1-2) | 83/620 (13·4%) | 85/636 (13·4%) | 57/364 (15·7%) | 59/373 (15·8%) | 49/284 (17·3%) |
| Upper Secondary(3-4) | 107/620 (17·3%) | 111/636 (17·5%) | 71/364 (19·5%) | 69/373 (18·5%) | 56/284 (19·7%) |
| Tertiary | 13/620 (2·1%) | 13/636 (2·0%) | 7/364 (1·9%) | 6/373 (1·6%) | 6/284 (2·1%) |
| Marital status ‡ | | | | | |
| Single | 106/638 (16·6%) | 107/656 (16·3%) | 66/375 (17·6%) | 61/383 (15·9%) | 53/293 (18·1%) |
| Married | 519/638 (81·3%) | 537/656 (81·9%) | 302/375 (80·5%) | 315/383 (82·2%) | 237/293 (80·9%) |
| Widowed | 3/638 (0·5%) | 2/656 (0·3%) | 2/375 (0·5%) | 3/383 (0·8%) | 2/293 (0·7%) |
| Divorced/Separated | 10/638 (1·6%) | 9/656 (1·4%) | 5/375 (1·3%) | 4/383 (1·0%) | 1/293 (0·3%) |
| Others | 0/638 (0·0%) | 1/656 (0·2%) | 0/375 (0·0%) | 0/383 (0·0%) | 0/293 (0·0%) |
| Income source ‡ | | | | | |
| None | 48/638 (7·5%) | 47/656 (7·2%) | 31/375 (8·3%) | 32/383 (8·4%) | 28/293 (9·6%) |
| Subsistence farming | 131/638 (20·5%) | 135/656 (20·6%) | 70/375 (18·7%) | 68/383 (17·8%) | 48/293 (16·4%) |
| Large scale farming | 2/638 (0·3%) | 2/656 (0·3%) | 1/375 (0·3%) | 1/383 (0·3%) | 1/293 (0·3%) |
| Employed | 79/638 (12·4%) | 80/656 (12·2%) | 47/375 (12·5%) | 50/383 (13·1%) | 37/293 (12·6%) |
| Casual work for wages | 212/638 (33·2%) | 226/656 (34·5%) | 118/375 (31·5%) | 124/383 (32·4%) | 87/293 (29·7%) |
| Business | 159/638 (24·9%) | 158/656 (24·1%) | 104/375 (27·7%) | 104/383 (27·2%) | 89/293 (30·4%) |
| Other | 7/638 (1·1%) | 8/656 (1·2%) | 4/375 (1·1%) | 4/383 (1·0%) | 3/293 (1·0%) |
| HIV positive ‡ | 90/634 (14·2%) | 92/652 (14·1%) | 55/372 (14·8%) | 59/379 (15·6%) | 44/291 (15·1%) |
| Malaria RDT positive ¶ | 9/627 (1·4%) | 11/647 (1·7%) | 4/369 (1·1%) | 3/378 (0·8%) | 3/291 (1·0%) |
| Re-screened post positive malaria RDT § | 212/641 (33·1%) | 214/659 (32·5%) | 129/377 (34·2%) | 134/385 (34·8%) | 104/295 (35·3%) |
| Venous Hb (g/dL) | 8·83 (1·21) | 8·83 (1·20) | 8·88 (1·22) | 8·90 (1·22) | 8·91 (1·25) |
| Anaemia status (venous) | | | | | |
| No | 30/638 (4·7%) | 28/655 (4·3%) | 20/375 (5·3%) | 21/381 (5·5%) | 18/294 (6·1%) |
| Mild | 74/638 (11·6%) | 77/655 (11·8%) | 46/375 (12·3%) | 46/381 (12·1%) | 37/294 (12·6%) |
| Moderate | 494/638 (77·4%) | 508/655 (77·6%) | 286/375 (76·3%) | 289/381 (75·9%) | 222/294 (75·5%) |
| Severe | 40/638 (6·3%) | 42/655 (6·4%) | 23/375 (6·1%) | 25/381 (6·6%) | 17/294 (5·8%) |
| Ferritin (µg/L) | 31·40 (10·50-79·30) | 28·90 (10·30-81·60) | 30·00 (9·30-80·70) | 27·90 (9·70-73·00) | 27·65 (9·20-66·20) |
| C-reactive protein (mg/L) | 5·35 (2·90-11·00) | 5·20 (2·90-11·40) | 5·50 (2·80-11·20) | 5·40 (2·80-11·20) | 5·65 (2·90-11·70) |
| Iron deficient ‖ | 245/626 (39·1%) | 263/643 (40·9%) | 158/375 (42·1%) | 164/383 (42·8%) | 132/294 (44·9%) |
| Iron deficient anaemia ‖ | 232/623 (37·2%) | 250/639 (39·1%) | 148/373 (39·7%) | 151/379 (39·8%) | 122/293 (41·6%) |
| Inflammation * | 330/626 (52·7%) | 334/643 (51·9%) | 203/375 (54·1%) | 203/383 (53·0%) | 157/294 (53·4%) |
| Anaemia and inflammation * | 313/623 (50·2%) | 318/639 (49·8%) | 191/373 (51·2%) | 190/379 (50·1%) | 145/293 (49·5%) |

No./total no. (%) are counts and percentages Plus-minus values are means ±SD. RDT denotes rapid diagnostic test (for *Plasmodium* parasitaemia), Hb haemoglobin, and HIV human immunodeficiency virus.

† Body mass index is the weight in kilograms divided by the square of the height in meters.
‡ Religion, Education, Marital status, Income source, parity, gravidity and HIV status were self-reported.
§ If women met the anaemia criteria but had a positive RDT, they were treated for malaria as per local protocols and deferred from enrolment. Women were invited to be re-screened – no earlier than seven days later – and be enrolled if they met the eligibility criteria (in these cases, parasitemia was assessed using microscopy due to persistence of antigen detection via RDT).
¶ Malaria RDT positive based on confirmatory RDT testing by lab personal on venous blood collected at enrolment
|| Iron deficient indicates serum ferritin<15ug/L or ferritin<30ug/L if C-reactive protein>5mg/L, and iron deficiency anaemia indicates Hb<11g/dL and serum ferritin<15µg/L or ferritin<30µg/L if C-reactive protein>5mg/L.
* Inflammation indicates C-reactive protein>5mg/L, and anaemia and inflammation indicates Hb<11.0g/dL and C-reactive protein>5mg/L.

**Supplementary Table 2. Baseline Characteristics for those with non-missing parasitemia by iron status**

| | Ferritin<15µg/L or ferritin<30µg/L if CRP>5mg/L | | Ferritin<15µg/L or ferritin<70µg/L if CRP>5mg/L | | |
|---|---|---|---|---|---|
| | **Iron deficient N = 245** | **Iron replete N = 381** | **Iron deficient N = 319** | **Iron replete N = 307** | **Total N = 626** |
| Age (years) | 24·5 (6·8) | 20·3 (5·0) | 23·4 (6·7) | 20·4 (5·0) | 22·0 (6·1) |
| Primigravid | 87/245 (35·5%) | 266/381 (69·8%) | 138/319 (43·3%) | 215/307 (70·0%) | 353/626 (56·4%) |
| Gestational age (weeks) | 22·4 (19·6-24·6) | 21·9 (19·6-24·0) | 22·4 (19·3-24·7) | 21·9 (19·6-24·0) | 22·0 (19·6-24·3) |
| Height (cm) | 155·5 (6·8) | 155·3 (6·4) | 155·3 (6·5) | 155·4 (6·6) | 155·4 (6·5) |
| Weight (kg) | 56·8 (8·8) | 54·3 (7·8) | 56·3 (8·7) | 54·3 (7·7) | 55·3 (8·3) |
| Body Mass Index (kg/m2) † | 23·5 (3·3) | 22·5 (2·8) | 23·3 (3·3) | 22·5 (2·7) | 22·9 (3·0) |
| Religion ‡ | | | | | |
| None | 2/244 (0·8%) | 0/379 (0·0%) | 2/318 (0·6%) | 0/305 (0·0%) | 2/623 (0·3%) |
| Christian | 174/244 (71·3%) | 252/379 (66·5%) | 225/318 (70·8%) | 201/305 (65·9%) | 426/623 (68·4%) |
| Muslim | 67/244 (27·5%) | 124/379 (32·7%) | 90/318 (28·3%) | 101/305 (33·1%) | 191/623 (30·7%) |
| Other | 1/244 (0·4%) | 3/379 (0·8%) | 1/318 (0·3%) | 3/305 (1·0%) | 4/623 (0·6%) |
| Education ‡ | | | | | |
| None | 2/237 (0·8%) | 0/369 (0·0%) | 2/310 (0·6%) | 0/296 (0·0%) | 2/606 (0·3%) |
| Lower Primary(1-5) | 62/237 (26·2%) | 80/369 (21·7%) | 74/310 (23·9%) | 68/296 (23·0%) | 142/606 (23·4%) |
| Upper Primary(6-8) | 87/237 (36·7%) | 177/369 (48·0%) | 124/310 (40·0%) | 140/296 (47·3%) | 264/606 (43·6%) |
| Lower Secondary(1-2) | 29/237 (12·2%) | 53/369 (14·4%) | 41/310 (13·2%) | 41/296 (13·9%) | 82/606 (13·5%) |
| Upper Secondary(3-4) | 48/237 (20·3%) | 56/369 (15·2%) | 60/310 (19·4%) | 44/296 (14·9%) | 104/606 (17·2%) |
| Tertiary | 9/237 (3·8%) | 3/369 (0·8%) | 9/310 (2·9%) | 3/296 (1·0%) | 12/606 (2·0%) |
| Income source ‡ | | | | | |
| None | 17/244 (7·0%) | 31/379 (8·2%) | 22/318 (6·9%) | 26/305 (8·5%) | 48/623 (7·7%) |
| Subsistence farming | 44/244 (18·0%) | 81/379 (21·4%) | 56/318 (17·6%) | 69/305 (22·6%) | 125/623 (20·1%) |
| Large scale farming | 1/244 (0·4%) | 1/379 (0·3%) | 2/318 (0·6%) | 0/305 (0·0%) | 2/623 (0·3%) |
| Employed | 38/244 (15·6%) | 39/379 (10·3%) | 45/318 (14·2%) | 32/305 (10·5%) | 77/623 (12·4%) |
| Casual work for wages | 72/244 (29·5%) | 136/379 (35·9%) | 96/318 (30·2%) | 112/305 (36·7%) | 208/623 (33·4%) |
| Business | 68/244 (27·9%) | 88/379 (23·2%) | 92/318 (28·9%) | 64/305 (21·0%) | 156/623 (25·0%) |
| Other | 4/244 (1·6%) | 3/379 (0·8%) | 5/318 (1·6%) | 2/305 (0·7%) | 7/623 (1·1%) |
| HIV positive ‡ | 60/242 (24·8%) | 28/377 (7·4%) | 65/315 (20·6%) | 23/304 (7·6%) | 88/619 (14·2%) |
| Malaria RDT positive ¶ | 2/240 (0·8%) | 7/372 (1·9%) | 2/312 (0·6%) | 7/300 (2·3%) | 9/612 (1·5%) |
| Re-screened post positive malaria RDT § | 20/245 (8·2%) | 190/381 (49·9%) | 56/319 (17·6%) | 154/307 (50·2%) | 210/626 (33·5%) |
| Venous Hb (g/dL) | 8·80 (1·26) | 8·84 (1·19) | 8·83 (1·26) | 8·82 (1·17) | 8·82 (1·21) |
| Anaemia status (venous) | | | | | |
| No | 11/243 (4·5%) | 19/380 (5·0%) | 16/317 (5·0%) | 14/306 (4·6%) | 30/623 (4·8%) |
| Mild | 25/243 (10·3%) | 46/380 (12·1%) | 33/317 (10·4%) | 38/306 (12·4%) | 71/623 (11·4%) |
| Moderate | 189/243 (77·8%) | 294/380 (77·4%) | 245/317 (77·3%) | 238/306 (77·8%) | 483/623 (77·5%) |
| Severe | 18/243 (7·4%) | 21/380 (5·5%) | 23/317 (7·3%) | 16/306 (5·2%) | 39/623 (6·3%) |
| Ferritin (µg/L) | 8·60 (6·20-12·40) | 62·20 (36·30-119·80) | 10·70 (6·80-26·80) | 81·50 (36·30-140·70) | 31·40 (10·50-79·30) |
| C-reactive protein (mg/L) | 6·60 (3·50-10·60) | 4·70 (2·50-11·20) | 7·00 (4·30-12·30) | 3·80 (2·10-8·40) | 5·35 (2·90-11·00) |

| | | | | | |
|---|---|---|---|---|---|
| Inflammation * | 152/245 (62·0%) | 178/381 (46·7%) | 226/319 (70·8%) | 104/307 (33·9%) | 330/626 (52·7%) |

No·/total no· (%) are counts and percentages Plus-minus values are means ±SD· RDT denotes rapid diagnostic test (for *Plasmodium* parasitaemia), Hb haemoglobin, and HIV human immunodeficiency virus.
† Body mass index is the weight in kilograms divided by the square of the height in meters.
‡ Religion, Education, Income source, gravidity and HIV status were self-reported.
§ If women met the anaemia criteria but had a positive RDT, they were treated for malaria as per local protocols and deferred from enrolment. Women were invited to be re-screened – no earlier than seven days later – and be enrolled if they met the eligibility criteria (in these cases, parasitemia was assessed using microscopy due to persistence of antigen detection via RDT).
¶ Malaria RDT positive based on confirmatory RDT testing by lab personal on venous blood collected at enrolment.
* Inflammation indicates C-reactive protein>5mg/L.

**Supplementary Table 3 · Baseline Characteristics excluding women who were malaria RDT-positive at screening, treated with AL and enrolled after microscopy-negative slide (re-screened) by iron status**

| | Ferritin<15µg/L or ferritin<30µg/L if CRP>5mg/L | | Ferritin<15µg/L or ferritin<70µg/L if CRP>5mg/L | | |
|---|---|---|---|---|---|
| | **Iron deficient N = 225** | **Iron replete N = 191** | **Iron deficient N = 263** | **Iron replete N = 153** | **Total N = 416** |
| Age (years) | 24·8 (6·8) | 21·4 (5·6) | 24·2 (6·8) | 21·5 (5·6) | 23·2 (6·5) |
| Primigravid | 76/225 (33·8%) | 114/191 (59·7%) | 100/263 (38·0%) | 90/153 (58·8%) | 190/416 (45·7%) |
| Gestational age (weeks) | 22·6 (19·1-24·7) | 22·1 (19·0-24·0) | 22·6 (19·0-24·7) | 21·9 (19·0-23·9) | 22·3 (19·0-24·4) |
| Height (cm) | 155·7 (7·0) | 155·6 (7·1) | 155·5 (6·9) | 155·8 (7·3) | 155·6 (7·0) |
| Weight (kg) | 57·1 (9·0) | 54·6 (8·2) | 56·8 (8·9) | 54·4 (8·2) | 55·9 (8·7) |
| Body Mass Index (kg/m2) † | 23·6 (3·4) | 22·5 (3·1) | 23·5 (3·4) | 22·4 (3·0) | 23·1 (3·3) |
| Religion ‡ | | | | | |
| None | 2/224 (0·9%) | 0/189 (0·0%) | 2/262 (0·8%) | 0/151 (0·0%) | 2/413 (0·5%) |
| Christian | 160/224 (71·4%) | 130/189 (68·8%) | 189/262 (72·1%) | 101/151 (66·9%) | 290/413 (70·2%) |
| Muslim | 61/224 (27·2%) | 58/189 (30·7%) | 70/262 (26·7%) | 49/151 (32·5%) | 119/413 (28·8%) |
| Other | 1/224 (0·4%) | 1/189 (0·5%) | 1/262 (0·4%) | 1/151 (0·7%) | 2/413 (0·5%) |
| Education ‡ | | | | | |
| None | 1/217 (0·5%) | 0/183 (0·0%) | 1/254 (0·4%) | 0/146 (0·0%) | 1/400 (0·2%) |
| Lower Primary(1-5) | 56/217 (25·8%) | 45/183 (24·6%) | 63/254 (24·8%) | 38/146 (26·0%) | 101/400 (25·2%) |
| Upper Primary(6-8) | 79/217 (36·4%) | 84/183 (45·9%) | 96/254 (37·8%) | 67/146 (45·9%) | 163/400 (40·8%) |
| Lower Secondary(1-2) | 26/217 (12·0%) | 27/183 (14·8%) | 32/254 (12·6%) | 21/146 (14·4%) | 53/400 (13·2%) |
| Upper Secondary(3-4) | 46/217 (21·2%) | 25/183 (13·7%) | 53/254 (20·9%) | 18/146 (12·3%) | 71/400 (17·8%) |
| Tertiary | 9/217 (4·1%) | 2/183 (1·1%) | 9/254 (3·5%) | 2/146 (1·4%) | 11/400 (2·8%) |
| Income source ‡ | | | | | |
| None | 17/224 (7·6%) | 13/189 (6·9%) | 17/262 (6·5%) | 13/151 (8·6%) | 30/413 (7·3%) |
| Subsistence farming | 38/224 (17·0%) | 43/189 (22·8%) | 43/262 (16·4%) | 38/151 (25·2%) | 81/413 (19·6%) |
| Large scale farming | 1/224 (0·4%) | 0/189 (0·0%) | 1/262 (0·4%) | 0/151 (0·0%) | 1/413 (0·2%) |
| Employed | 35/224 (15·6%) | 22/189 (11·6%) | 41/262 (15·6%) | 16/151 (10·6%) | 57/413 (13·8%) |
| Casual work for wages | 64/224 (28·6%) | 57/189 (30·2%) | 77/262 (29·4%) | 44/151 (29·1%) | 121/413 (29·3%) |
| Business | 65/224 (29·0%) | 52/189 (27·5%) | 79/262 (30·2%) | 38/151 (25·2%) | 117/413 (28·3%) |
| Other | 4/224 (1·8%) | 2/189 (1·1%) | 4/262 (1·5%) | 2/151 (1·3%) | 6/413 (1·5%) |
| HIV positive ‡ | 57/222 (25·7%) | 21/189 (11·1%) | 61/260 (23·5%) | 17/151 (11·3%) | 78/411 (19·0%) |
| Malaria RDT positive ¶ | 1/220 (0·5%) | 3/186 (1·6%) | 1/256 (0·4%) | 3/150 (2·0%) | 4/406 (1·0%) |
| Re-screened post positive malaria RDT § | 0/225 (0·0%) | 0/191 (0·0%) | 0/263 (0·0%) | 0/153 (0·0%) | 0/416 (0·0%) |
| Venous Hb (g/dL) | 8·79 (1·25) | 8·94 (1·25) | 8·83 (1·25) | 8·91 (1·26) | 8·86 (1·25) |
| Anaemia status (venous) | | | | | |
| No | 10/223 (4·5%) | 15/190 (7·9%) | 14/261 (5·4%) | 11/152 (7·2%) | 25/413 (6·1%) |
| Mild | 23/223 (10·3%) | 26/190 (13·7%) | 28/261 (10·7%) | 21/152 (13·8%) | 49/413 (11·9%) |
| Moderate | 173/223 (77·6%) | 137/190 (72·1%) | 200/261 (76·6%) | 110/152 (72·4%) | 310/413 (75·1%) |
| Severe | 17/223 (7·6%) | 12/190 (6·3%) | 19/261 (7·3%) | 10/152 (6·6%) | 29/413 (7·0%) |
| Ferritin (µ/L) | 8·10 (6·20-11·90) | 59·20 (33·70-123·10) | 9·30 (6·40-15·70) | 77·70 (27·10-139·80) | 16·35 (7·85-52·55) |

| | | | | | |
|---|---|---|---|---|---|
| C-reactive protein (mg/L) | 6·40 (3·40-10·50) | 5·00 (2·70-11·40) | 6·90 (3·80-11·70) | 3·90 (2·30-9·00) | 5·65 (3·10-11·00) |
| Inflammation * | 138/225 (61·3%) | 90/191 (47·1%) | 176/263 (66·9%) | 52/153 (34·0%) | 228/416 (54·8%) |

No·/total no· (%) are counts and percentages Plus-minus values are means ±SD· RDT denotes rapid diagnostic test (for *Plasmodium* parasitaemia), Hb haemoglobin, and HIV human immunodeficiency virus.

† Body mass index is the weight in kilograms divided by the square of the height in meters.

‡ Religion, Education, Income source, gravidity and HIV status were self-reported.

§ If women met the anaemia criteria but had a positive RDT, they were treated for malaria as per local protocols and deferred from enrolment. Women were invited to be re-screened – no earlier than seven days later – and be enrolled if they met the eligibility criteria (in these cases, parasitemia was assessed using microscopy due to persistence of antigen detection via RDT).

¶ Malaria RDT positive based on confirmatory RDT testing by lab personal on venous blood collected at enrolment

* Inflammation indicates C-reactive protein>5mg/L.

**Supplementary Table 4. Comparison by WHO definition of iron status at baseline and qPCR positivity**

| | **Iron deficient at baseline** | | **Iron replete at baseline** | | **Unadjusted*** | | **Adjusted*** | |
|---|---|---|---|---|---|---|---|---|
| **PCR positive** | **N** | **n/N (%)** | **N** | **n/N (%)** | **Effect size (95% CI)** | **p-value** | **Effect size (95% CI)** | **p-value** |
| Baseline* | 319 | 92/319 (28·8) | 307 | 186/307 (60·6) | 0·48 (0·37, 0·611) | <0·0001 | 0·62 (0·47, 0·81) | 0·0006 |
| 28 days post treatment+ | 336 | 50/336(14.9) | 307 | 65/307(21.2) | 0·71(0·49, 1.02) | 0.064 | 0·87 (0·56, 1.34) | 0.52 |
| Excluding those malaria RDT-positive at screening, treated with AL and enrolled after microscopy-negative slide (re-screened) | | | | | | | | |
| Baseline* | 263 | 61/263 (23.2) | 153 | 88/153 (54.6) | 0·40 (0·29, 0·56) | <0·0001 | 0·49 (0·35, 0·70) | <0·0001 |
| 28 days post treatment+ | 279 | 39/279 (14.0) | 152 | 35/152 (23.0) | 0·61 (0·38, 0·96) | 0.032 | 0·75 (0·45, 1·25) | 0·27 |

Iron deficient indicates serum ferritin <15 µg/L or ferritin <70 µg/L if C-reactive protein >5 mg/L

*Unadjusted: N/A;

Adjusted: gestational age at baseline, gravidity status, HIV, rescreened + AL treatment, maternal age, education, income source, and religion

+Unadjusted: treatment group:

Adjusted: treatment group, gestational age at baseline, gravidity status, HIV, rescreened + AL treatment, maternal age, education, income source, and religion

*Poisson regression model with robust error variance was fit

**Supplementary Table 5. Prevalence of qPCR-positivity at 28 days post treatment in qPCR-negative women at baseline by iron status**

| | Iron deficient at base | | Iron replete at base | | Unadjusted * | | Adjusted* | |
|---|---|---|---|---|---|---|---|---|
| **PCR positive** | **N** | **n/N (%)** | **N** | **n/N (%)** | **Effect size (95% CI)** | **p-value** | **Effect size (95% CI)** | **p-value** |
| 28 days post treatment+ | 215 | 16/215 (7·4) | 177 | 20/177 (11·3) | 0·66 (0·34, 1·27) | 0·21 | 0·95 (0·41, 2·19) | 0·91 |

Unadjusted: N/A;
adjusted: Gestational age at baseline, gravidity status, HIV, rescreened + AL treated, maternal age, education, income source, religion
+Unadjusted: treatment group:
+Adjusted: treatment group, gestational age at visit 2, gravidity, HIV, rescreened + treated, maternal age, education, income source, religion
*Poisson regression model with robust error variance was fit

**Supplementary Table 6. Parasite densities at 28 days post treatment in qPCR-negative women at baseline, by iron status at baseline**

| | **Iron deficient at baseline** | **Iron replete at baseline** | **Overall** |
|---|---|---|---|
| | N=16 | N=20 | N=36 |
| parasites per mL of blood | 3·27 (0·70-439·66) | 19·60 (0·33-558·50) | 4·48 (0·35-522·32) |
| $Log_e$ parasites per mL of blood | 1·18 (-0·49-6·00) | 2·97 (-1·11-6·30) | 1·48 (-1·04-6·24) |

Values represent the median and IQR.

Due to small sample sizes and violation of assumptions, only summary data are presented for the continuous parasite densities without statistical analysis comparing groups.

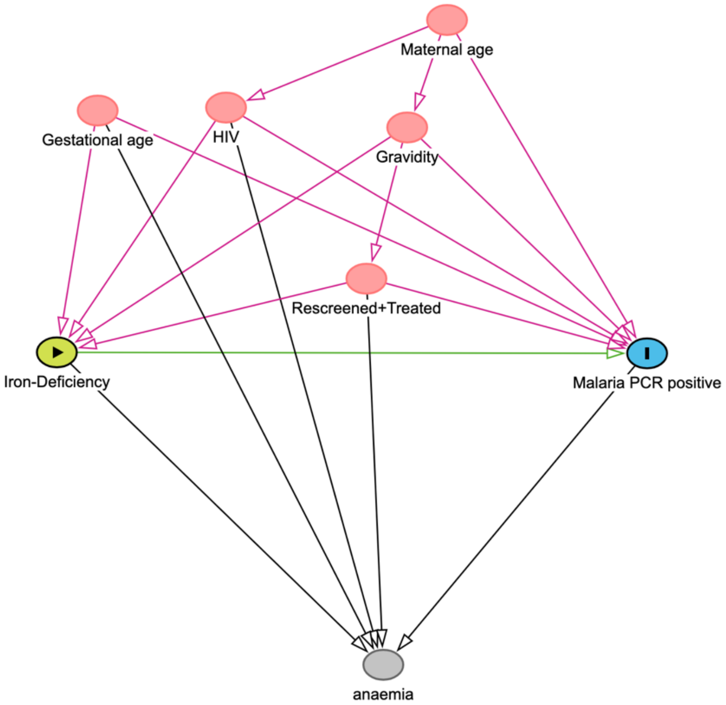


**Supplementary figure 1. Directed Acyclic Graph of the association between Iron-deficiency and *Plasmodium* qPCR-positivity at trial baseline**. Arrows indicate the direction of the assumed causal influence. The primary exposure of interest is iron-deficiency status at baseline (green node) and the outcome is *Plasmodium* infection detected by qPCR at baseline (blue node). Measured variables that influence both the exposure and the outcome are shown (red nodes). Variables not adjusted for in the main analysis are indicated in grey. This DAG was constructed in DAGitty (http://www.dagitty.net) based on existing literature and subject-matter knowledge to guide variable selection for the minimal adjustment set to obtain the total effect and identify potential sources of bias.

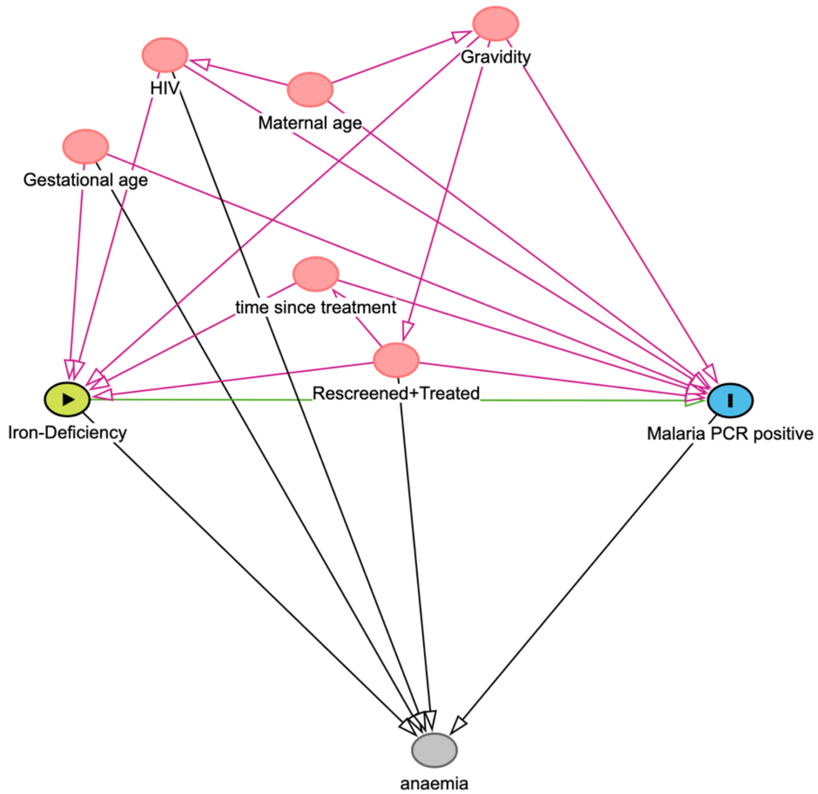


**Supplementary figure 2. Directed Acyclic Graph of the association between Iron-deficiency and *Plasmodium* qPCR-positivity at 28 days post treatment**. Arrows indicate the direction of the assumed causal influence. The primary exposure of interest is iron-deficiency status at baseline (green node) and the outcome is *Plasmodium* infection detected by qPCR at 28 days post treatment (blue node). Measured variables that influence both the exposure and the outcome are shown (red node). Variables not adjusted for in the main analysis are indicated in grey. This DAG was constructed in DAGitty (http://www.dagitty.net) based on existing literature and subject-matter knowledge to guide variable selection for the minimal adjustment set to obtain the total effect and identify potential sources of bias.

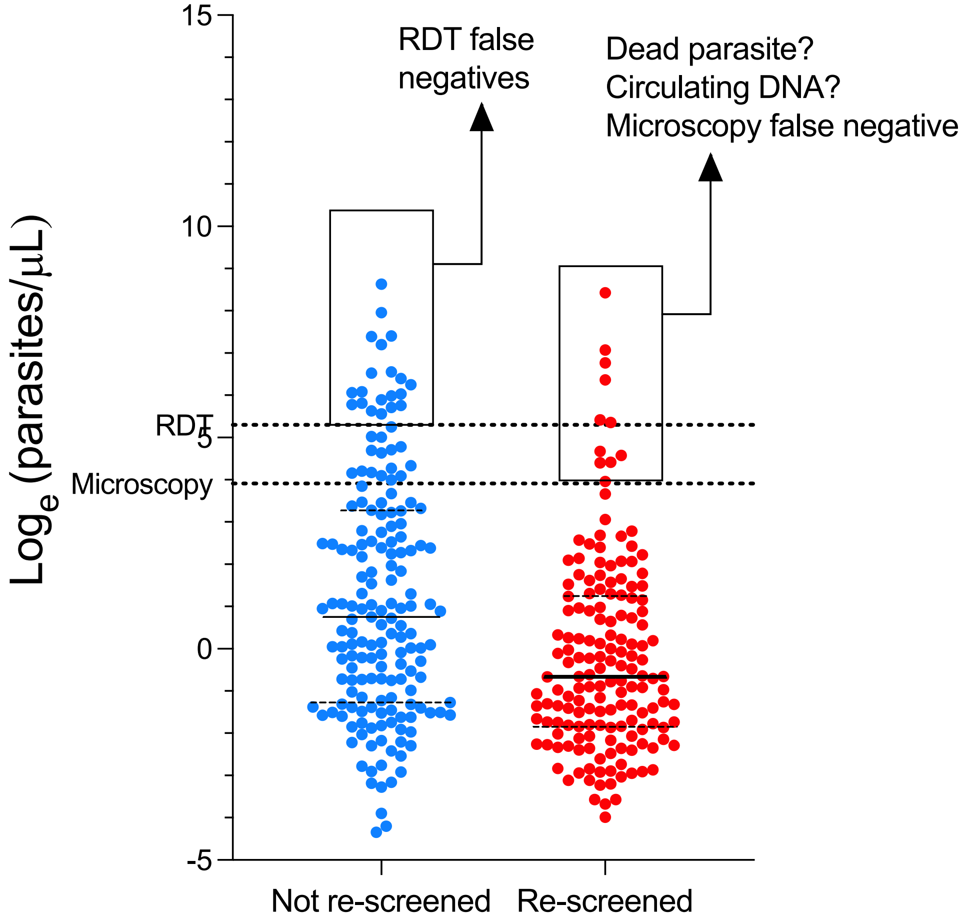


**Supplementary figure 3. *P falciparum* density by screening status.** Parasite density as measured by ultrasensitive qPCR (log$_e$ parasites/mL) in women at baseline, comparing those who entered the study directly (not re-screened (blue)) with those who entered the study after being initially RDT-positive at the time of screening, treated with artemether-lumefantrine, and allowed to be re-screened and enrolled if they had a negative microscopy slide (Re-screened (red)). Dotted lines represent the theoretical limits of detection for malaria RDTs (200 parasites/μL) and microscopy slides (50 parasites/μL).

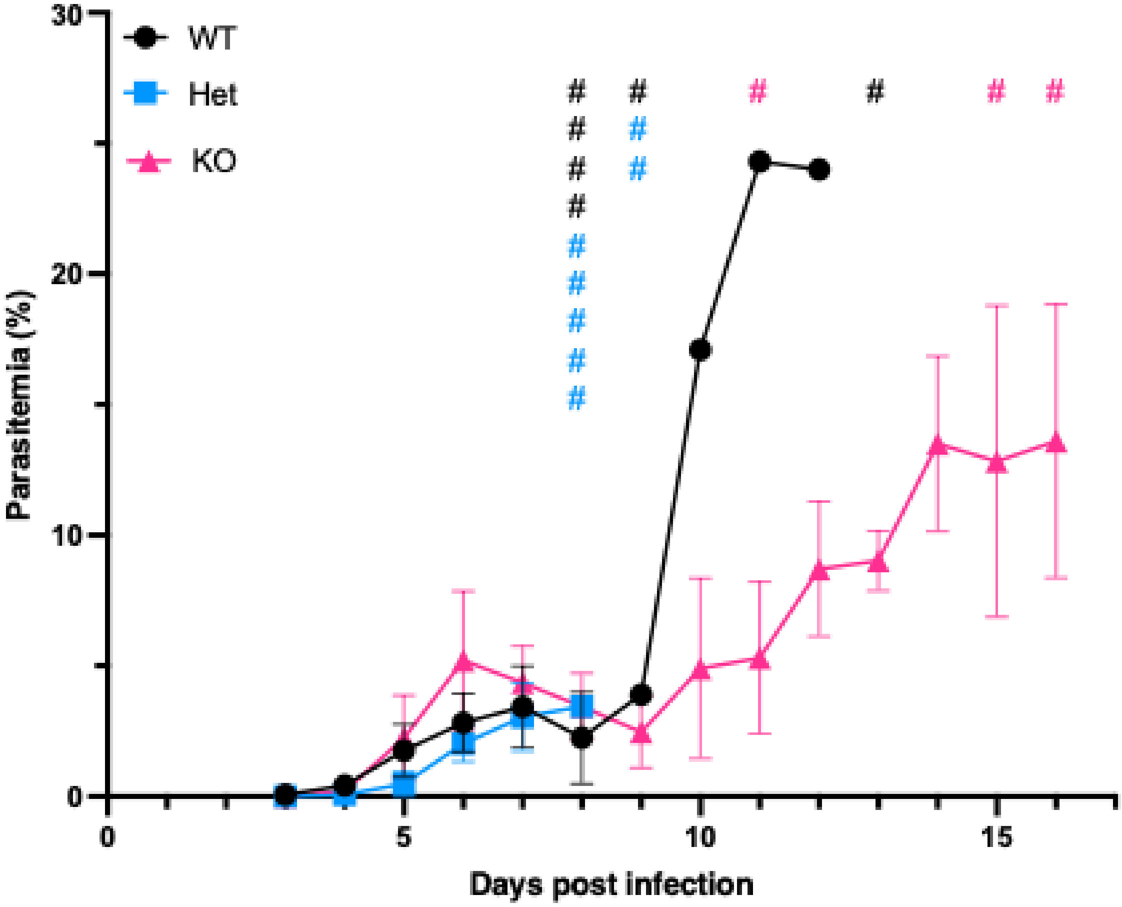


**Supplementary figure 4. Blood-stage parasitaemia up to day 16 post infection.** *P. berghei* blood-stage parasitaemia in C57BL/6 wild-type (WT), *Tmprss6*-heterozygous (Het), and *Tmprss*6-knockout (KO) mice. Data represented as mean ± SD, n=6-7 mice per group. # = euthanasia.
